# Marine *Gammaproteobacteria* use SusC/D-like proteins in fructan utilization

**DOI:** 10.1101/2024.09.11.612387

**Authors:** Marie-Katherin Zühlke, Alexandra Bahr, Daniel Bartosik, Vipul Solanki, Michelle Teune, Norma Welsch, Frank Unfried, Tristan Barbeyron, Elizabeth Ficko-Blean, Paula Schoppmeier, Laurie Schiller, Lionel Cladière, Alexandra Jeudy, Anne Susemihl, Fabian Hartmann, Diane Jouanneau, Murielle Jam, Matthias Höhne, Mihaela Delcea, Uwe T. Bornscheuer, Dörte Becher, Jan-Hendrik Hehemann, Mirjam Czjzek, Thomas Schweder

**Author notes:** corresponding authors: Marie-Katherin Zühlke,; Thomas Schweder.

## Abstract

Fructans are ubiquitous in terrestrial ecosystems, however, these glycans are unexplored in the marine environment. We have discovered that the Antarctic gammaproteobacterium *Pseudoalteromonas distincta* is highly adapted to the degradation of fructose-containing substrates. This is enabled by proteins encoded in several genomic regions, including a fructan polysaccharide utilization locus (PUL). In addition to a glycoside hydrolase from family 32 (GH32), the fructan PUL encodes two proteins that have been described as specific for *Bacteroidota* and were previously unknown for *Gammaproteobacteria*: a glycan-binding SusD-like protein and a SusC-like TonB-dependent transporter (TBDT), which work as a complex in glycan import. Proteome analyses and biochemistry results suggest that the SusC/D-like proteins of *P. distincta* shuttle small-sized inulin-type fructans directly into the cell, where they are degraded by a periplasmic exo-active GH32. A SusD-like protein could provide a competitive adavantage in the absence of extracelluar endo-active inulinases. Comparative genomics identified further SusC/D-like proteins in *Gammaproteobacteria*, most of which are co-encoded with GH32s, indicative of fructan PULs, and are frequently associated with the marine habitat. Our study thus shows the first known exception to the paradigm that only *Bacteroidota* use SusC/D-like proteins. It further suggests that fructans contribute to the marine glycan pool.

## Introduction

We are only beginning to understand the diversity of glycan structures that marine algae - but also fungi and bacteria - produce and how such glycans contribute to carbon flux in the ocean. Marine *Bacteroidota* and *Gammaproteobacteria* are specialists in the degradation of such oceanic glycans and have evolved unique glycan utilization strategies.

In the *Bacteroidota* phylum, gene clusters encoding glycan-disassembling carbohydrate-active enzymes (CAZymes), such as glycoside hydrolases (GHs), and glycan-transporting SusC/D-like complexes are referred to as polysaccharide utilization loci (PULs), where ‘sus’ refers to homologs of the starch utilization system (sus) of the human intestinal *Bacteroides thetaiotaomicron* (1, 2). SusC/D-like proteins are structurally designed to function as a complex, in which the outer membrane-tethered SusD-like lipoprotein binds and transmits the oligosaccharides released by extracellular CAZymes to the SusC-like TonB-dependent receptors or transducers (referred herein as transporters, TBDTs) using the ‘pedal-bin’ mechanism (3). Compared to ‘classical’ TBDTs, SusC-like TBDTs are larger to accommodate the SusD-like protein (4). In contrast, carbohydrate utilization clusters of *Gammaproteobacteria* (phylum *Pseudomonadota*), often referred to as PULs as well, lack SusD-like proteins and the “SusC-like” specific characteristics of TBDTs. Surprisingly, we have discovered a predicted SusC-like TBDT/SusD-like protein pair encoded in a PUL of the Antarctic gammaproteobacterium *Pseudoalteromonas distincta* ANT/505. In addition to these *Bacteroidota*-specific proteins, the PUL encodes a GH32 which suggests fructan utilization (5), a substrate which is unexplored in the marine ecosystem. Fructans are polyfructose chains, in which fructosyl units are β(2,1)- or β(2,6)-linked to sucrose, referred to as inulin or levan, respectively. In terrestrial ecosystems, where fructan production is widespread (6), fructans serve as storage compounds or osmolytes in plants and are frequently found as exopolysaccharides (EPS) in Gram-positive and Gram-negative bacteria. In bacteria, levan and inulin are produced by GH68s (6).

The ability to degrade fructans is widespread among terrestrial bacteria (7) and has been thoroughly investigated in *Bacteroidota* of the human intestine (8–11). For the marine counterpart, inulinase activity of marine fungi and bacteria (12–15) or the presence of GH32-containing PULs in metagenome-assembled genomes of marine bacteria (16, 17) have been reported, but in-depth biochemical and structural analyses of the corresponding CAZymes, as in the case of the *Thermotoga maritima* GH32 (15, 18), are rare. Moreover, little is known about fructan utilization pathways in general.

In this study, we describe the first marine utilization pathway for fructans in the Antarctic gammaproteobacterium *P. distincta*. Our results show that specialized marine *Gammaproteobacteria* degrade fructans using PULs that encode SusC/D-like transport complexes and GH32s. To the best of our knowledge, this is the first report of SusC/D-like proteins in *Gammaproteobacteria*.

## Materials and Methods

### Carbohydrates

Plant-derived levan (Timothy grass) and fructo-oligosaccharides (FOS_INU_, DP2-8) from inulin (chicory) were purchased from Megazyme/NEOGEN, inulin (chicory) from Alfa Aesar (VWR) and bacterial levan (*Erwinia herbicola*), pullulan and maltotriose from Sigma Aldrich (Merck).

### Cultivation of P. distincta

Cells of overnight cultures in marine broth 2216 were pelleted and washed with an artificial seawater medium (MPM) (19). This suspension was then used to inoculate main cultures of *P. distincta* in MPM supplemented with 0.2% of the respective substrates to an OD_600nm_ of 0.1. Inoculated MPM medium without any carbon source served as a control. Cells were cultivated at 10 °C and 150 rpm.

### Proteome analysis

Cells were harvested by centrifugation (15 min, 9,500 x *g*, 4 °C) in the late exponential growth phase. For whole-cell proteomes, cell pellets were resuspended in TE-buffer (10 mM Tris, 10 mM EDTA) and cells were lysed via sonication (4 cycles of 25 s, 30% power, pulse 5, Bandelin Sonopuls HD 2070 ultrasonic homogenizer). Cell debris was removed by centrifugation. Intracellular soluble fractions, outer membranes and outer membrane vesicles were prepared as described before (20–22). Proteins were extracted from the supernatant using StrataClean beads according to published protocols (23, 24), referred to as the extracellular proteome. Here, 20 µL of bead solution was used to extract 20-25 µg of protein. Otherwise, the Pierce BCA assay determined protein concentration and 25 µg of protein was separated by 1D-SDS-PAGE. Lanes were cut into 10 equal pieces and proteins were digested with trypsin as described before (24). Peptides were subjected to reversed phase chromatography and analyzed using an online-coupled LTQ Orbitrap (whole-cell proteomes) or LTQ XL Orbitrap (subcellular fractions) mass spectrometer equipped with a nanoelectrospray ion source (Thermo Fisher Scientific Inc., Waltham, MA, USA). Data analysis was performed using MaxQuant v. 1.6.0.16 with the integrated Andromeda search engine (25). The false discovery rate (FDR) was set to 1% on peptide and protein level and match between runs was enabled (for subcellular protein fractions only between replicates). %riBAQ values (relative intensity-based absolute quantification giving the relative protein abundance in the sample) were calculated to determine protein abundance per sample. Proteins that were quantified in at least two out of three replicates were considered for calculation of mean values.

For statistical analysis in the case of whole-cell proteomes, the data set was filtered for at least one valid value per protein and missing values were imputed with a small constant. Values were then log-transformed and row-wise z-score transformation was used to analyze differential production of proteins. Significant differences were determined by one-way analysis of variance (ANOVA) followed by a Tukey’s honestly significant difference (HSD) post hoc test (FDR 5%) in Perseus v. 2.1.0.0 (26, 27) using the filtered, imputed and log-transformed abundance values of proteins. The mass spectrometry proteomics data have been deposited to the ProteomeXchange Consortium via the PRIDE (28, 29) partner repository, see Data Availability Statement. PSORTb v. 3.0 (30), DeepLocPro v. 1.0 (31), SignalP v. 6.0 (32) and DeepSig (33) were used to evaluate suggested subcellular localization of proteins.

### DNA cloning

The Gibson assembly strategy (34) was used to clone genes into the pET28a vector with primers listed in Table S1. Primers were designed to produce *P. distincta* proteins as recombinant N-terminal His_6_-tagged constructs, removing predicted signal peptides from sequences (32, 33). The NEBuilder^®^ HiFi DNA Assembly Master Mix (NEB) was used for assembly following the manufacturer’s protocol. Sequencing (Eurofins) confirmed sequence identity. For protein production, plasmids were transformed into competent *Escherichia coli* BL21(DE3) cells. Since PdGH32_DIS_ was not active using the predicted open reading frame (ORF), the corresponding gene was re-cloned using an alternative ORF (for details see Table S2) resulting into active CAZyme.

### Protein production & purification

*E. coli* BL21(DE3) clones, harbouring the corresponding plasmids, were grown in lysogeny broth (LB) containing 30 µg mL^-1^ kanamycin at 37 °C and 120 rpm. Medium was cooled down before isopropyl β-D-1-thiogalactopyranoside (IPTG) induction (0.3 or 1 mM) and cells were grown overnight at 20 °C at 120 rpm. In the case of CAZyme activity analysis, proteins were produced using the ZYP5052 autoinduction medium (35), except for the re-cloned PdGH32_DIS_, where the Enpresso® B medium (Enpresso GmbH, Berlin, Germany) was used according to the manufacturer’s instructions. Cells were harvested by centrifugation (e.g., 1 L culture: 35 min, 13,881 x *g*, 4 °C) and pellets were stored at -20 °C until further treatment. A chemical lysis protocol (36) was used to disrupt the cells (resuspension buffer: 50 mM Tris-HCl pH 8.0, 25% sucrose, ∼10 mg lysozyme; lysis buffer: 20 mM Tris-HCl pH 7.5, 100 mM NaCl, 1% sodium deoxycholate, 1% Triton-X). For CAZyme activity tests, cells were lysed via ultrasonication on ice (Bandelin Sonoplus HD2200, KE76 sonotrode; 2 x 3 min, 1 min pause, 50% power, 0.5 pulse) in buffer A (300 mM NaCl, 20 mM Tris-HCl pH 8.0, 20 mM imidazole).

For protein purification, 1 ml or 5 mL HisTrap HP columns were used, equilibrated in buffer A (300-500 mM NaCl, 20 mM Tris-HCl pH 8.0, 20 mM imidazole) and charged with 0.1 M NiSO_4_. The proteins were eluted using a linear gradient of 0-100% buffer B (300-500 mM NaCl, 20 mM Tris-HCl pH 8.0, 500 mM imidazole) within 20-60 min. If necessary, the purity of the sample was increased by size-exclusion chromatography (SEC) using a Superdex 200 16/60 column (Cytiva) in buffer C (CAZyme activity: 25 mM Tris-HCl pH 8.0, 50 mM NaCl, otherwise: 20 mM Tris-HCl pH 8.0, 200-250 mM NaCl). Otherwise, samples were desalted using a HiPrep 26/10 column and buffer C. Protein concentration was determined using a Nanodrop Spectrophotometer using the molecular weight and calculated extinction coefficient (37). If necessary, proteins were concentrated at 1,890 x *g* and 4 °C using Amicon Ultra centrifugal filter units (Merck) or using a stirred cell ultrafiltration device (10 kDa cut-off).

### Reducing sugar assay (RSA)

The degradation of different oligo- and polysaccharides by CAZymes was carried out using final concentrations of 2 mg mL^-1^ substrate and 30 µg mL^-1^ of CAZyme in 1 x PBS, pH 6.0 (120 µg mL^-1^ in case of the re-cloned PdGH32_DIS_). The enzymatic reactions were incubated for 24 h at room temperature. For the quantification of reducing ends released upon degradation, 20 µL of the reaction solutions were mixed with 20 µL DNS-reagent (0.4 M NaOH, 30% potassium sodium tartrate, 1% 3,5-dinitrosalicylic acid; DNS) and incubated at 100 °C for 15 min. dH_2_0 (180 µL) was added and 200 µL of the mixtures were transferred to a microtiter plate. The absorption at 540 nm was measured using a Tecan infinite 200 pro.

### Fluorophore-assisted carbohydrate electrophoresis (FACE)

The enzymatic reactions (10 µL) were lyophilized, resuspended in ANTS (0.1 M in DMSO, 4 µL, Invitrogen™) and NaCNBH_3_ (1 M in DMSO, 4 µL) and incubated at 37 °C overnight. The labeled samples were mixed with loading dye (62 mM Tris pH 6.8, 0.014% bromophenol blue, 10% glycerol, 8 µL) and separated at 400 V on a 30% polyacrylamide running gel with a 4% stacking gel for about 1h under consistent cooling. Imaging was performed using a 312 nm UV table.

### Isothermal titration calorimetry (ITC)

PdSusD_PUL_ was dialysed against 50 mM Tris pH 8.0, 100 mM NaCl at 4 °C for at least 48 h. ITC analysis was performed using a MicroCal^TM^ ITC 200 machine. The dialyses buffer was used to dissolve the substrates and to wash the ITC sample cell. All samples were centrifuged before analysis. Injections with 20 mM of inulin (calculated as DP17), 5 mM of plant-derived levan, 7.5 µM of bacterial levan and 20 mM of sucrose or fructose detected no signal. Since little heat was released during titrations at a concentration of 31.7 mM FOS_INU_ (calculated as DP5) and due to good solubility at high concentrations, we tested even higher concentrations of FOS_INU_: 63.5 mM of FOS_INU_ were injected into 160.97 µM of PdSusD_PUL_, which was performed in triplicate. The following settings were selected: cell temperature 20 °C (293.15 K), reference power 10 µCal/s, stirring speed 500 rpm, filter period 1 sec, injection spacing 200 sec, 2 µL injection volume (first injection 0.3 µL), 18 injections in total. As a control, FOS_INU_ was titrated into buffer and buffer into buffer. Due to the fact that the low affinity resulted in a C-value of 0.01 (38) and to the best of our knowledge SusD-like proteins bind in a 1:1 ratio, the n-value was fixed to 1. MICROCAL ORIGIN v.7 was used to analyze the data applying a single-site binding model.

### Protein structure prediction and analysis

AlphaFold2-predicted structures were downloaded from the AlphaFold Protein Structure DB (39), which is integrated in the Uniprot knowledgebase (40) and visualized using PyMOL Molecular Graphics System v2.5.4 (Schrödinger, LLC.).

### Phylogenetic analysis

A BLAST (41) of PdSusD_PUL_ against the UniProtKB and Reference Proteomes retrieved 1,750 unique sequences. Additional SusD-like protein sequences encoded in fructan PULs from *Bacteroidota* of the human intestine (with suspected or known specificity for inulin or levan (3, 8, 11)) as well as SusD-like protein-coding sequences from PULs of *Zobellia galactanivorans* Dsij^T^ with a substrate specificity other than fructan (42, 43) were added to the dataset. Thus, in total, 1776 sequences were aligned using MAFFT v. 7 (44) using the L-INS-i algorithm. The obtained multiple sequence alignment was visualized and processed in Jalview v. 11.0 (45). Short sequences, signal peptides and non-aligned regions were removed. Sequences were filtered at 90% identity and finally, 1055 sequences with 410 positions were used for the phylogeny. The phylogenetic tree was built by the maximum likelihood method (46) using RAxML v. 8.2.4 (47). The best evolutive model was selected submitting the alignment to IQ-TREE web server (http://iqtree.cibiv.univie.ac.at) (48). The LG matrix as evolutive model (49) and a discrete Gamma distribution to model evolutionary rate differences among sites (4 categories) were used for the calculation. The rate variation model allowed for some sites to be evolutionarily invariable. The reliability of the trees was tested by bootstrap analysis using 1000 resamplings of the dataset (50). The tree was visualized using iTOL v. 5 (51) and midpoint rooted.

### Co-occurrence analysis

Gammaproteobacterial genomes of the unified genome catalog of marine prokaryotes (UGCMP) (52) and the Genomes from Earth’s Microbiomes (GEM) catalog (53) were dereplicated using dRep (v3.4.2, flags: -l 0 -comp 50 -con 10 -pa 0.95 --checkM_method taxonomy_wf) (54). Resulting 4,681 genomes were reannotated using prokka (v1.14.6) (55) and screened for SusD-like homologs using hmmsearch (v3.3.2, gathering threshold) (56) with PFAM model PF07980.15 (SusD_RagB), PF12741.11 (SusD-like), PF12771.11 (SusD-like_2) and PF14322.10 (SusD-like_3) (57).

In order to examine polysaccharide utilization-related functions around *susD*-like genes, relevant functions were predicted using a five-gene sliding window. For TBDT annotation, sequences were compared to PF13715.11 (CarboxypepD_reg-like domain), PF07715.18 (Plug) and PF00593.27 (TonB_dep_Rec) using hmmsearch (gathering threshold). Sequences containing PF07715.18 and PF00593.27 were kept for further analysis. TBDT also fitting the gathering threshold for TIGRFAM profile TIGR04056 (OMP_RagA_SusC) (58) were classified as SusC-like proteins. CAZyme families were assigned with hmmscan vs. dbCAN-HMMdb-V12 and dbCAN-sub as well as diamond blastp (v2.1.1.155, e-value cut-off 1E-102) against CAZyDB.07262023, all provided by dbCAN (59). Results were then filtered using the hmmscan-parser.sh script with an e-value cutoff of 1E-15 and a minimum coverage of 0.35. CAZyme families predicted by at least two tools were kept for further analysis, not considering GT and AA assignments. Results were visualized with the ComplexUpset R package (v1.3.5) (60).

Furthermore, the 4,681 representative gammaproteobacterial genomes were searched for GH32 family encoding genes with hmmsearch (v3.3.2, option settings: -E 1 --domE 1 --incE 0.01 --incdomE 0.03) using GH32 HMM profiles from dbCAN-HMMdb-V12 and dbCAN-sub. Hits were further assigned to a final CAZyme family as described above. Their genetic neighborhood 5 genes up- and downstream was reconstructed similar to the predictions mentioned above, including hmmscan (gathering threshold) against the full Pfam-A database (as of 20240508).

## Results

### Pseudoalteromonas distincta contains several genomic regions to utilize fructose-containing substrates

The *P. distincta* fructan PUL classifies as a *Bacteroidota* PUL. Upstream of a GH32 (PdGH32_PUL_), the PUL encodes a SusC-like TBDT (PdSusC_PUL_) and a SusD-like protein (PdSusD_PUL_) (Fig. 1A). In addition to the receptor/β-barrel domain PF00593 and plug domain PF07715 that identify ‘classical’ TBDTs, the SusC-like subclade SusC/RagA model TIGR04056 was identified for the transport protein PdSusC_PUL_ (SusC/D homologs are named RagA/B in the human oral *Porphyromonas gingivalis* and are specific for oligopeptides (61)). In line with this, the Pfam family for SusD/RagB homologs PF07980 was identified for the co-encoded putative glycan-binding protein PdSusD_PUL_, which contains a lipoprotein signal peptide.

Additional genomic regions are presumably also related to the utilization of fructans or fructose-containing substrates by *P. distincta*: a distal GH32 (PdGH32_DIS_) and GH68 (PdGH68_DIS_), as well as a PUL that encodes proteins to cleave sucrose (GH13 subfamily 18, GH13_18) and to metabolize fructose (fructokinase) (Fig. 1A). However, the PUL also encodes a GH16_3, which could indicate β-glucan use (62). It is unknown whether this genetic region is actually one PUL dedicated to a common substrate or whether it is two PULs.

**Fig. 1.**
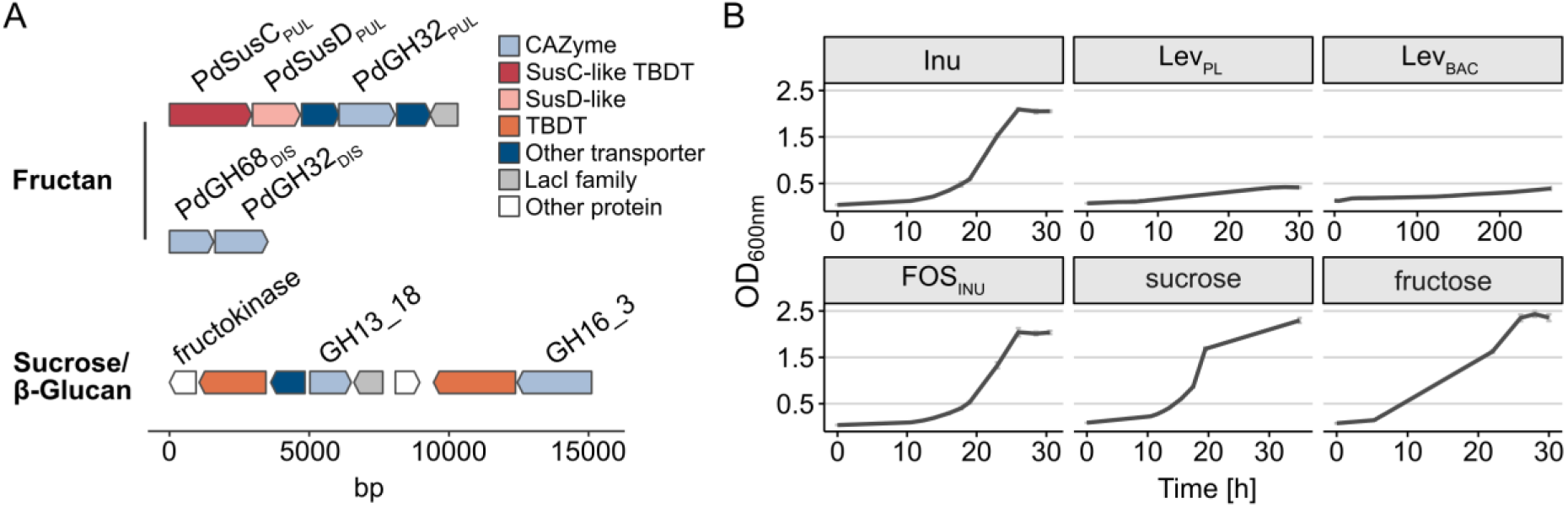
The gammaproteobacterium *P. distincta* is spezialized in the degradation of fructose-containing substrates. (**A**) The genomic context of two GH32-encoding genes, one in a fructan PUL (PdGH32PUL) and one distal to the PUL (PdGH32DIS), is shown. In addition, the *P. distincta* contains a genomic region likely specific for sucrose (GH13_18) and β-glucans (GH16_3 and co-encoded TBDT). The cluster might be even longer, see Fig. 2. (**B**) Growth of *P. distincta* with fructans, FOSINU, sucrose and fructose. There is a different time scale for bacterial levan. Controls without the addition of a carbon source are shown in Fig. S1. Inu: inulin, LevPL: plant-derived levan, LevBAC: bacterial levan, FOSINU: fructo-oligosaccharides from inulin.

Using fructans of terrestrial origin, we confirmed that *P. distincta* grows on inulin, oligosaccharides from inulin (FOS_INU_), sucrose and fructose, but reaches only low optical densities on plant-derived levan and hardly grows on bacterial levan (Fig. 1B, for a control without substrate, see Fig. S1).

### Inulin and levan stimulate protein production of fructan PUL-encoded proteins

The relative abundance of proteins (%riBAQ) encoded in the different genomic regions associated with the use of fructans was increased in cells grown with inulin, FOS_INU_ and both levans compared to cells grown with α-glucans (Fig. 2). Fructan PUL-encoded proteins were abundant in whole cell proteomes of cells cultivated with inulin, FOS_INU_ and plant-derived levan, but also the bacterial levan (Fig. S2, Supplementary Dataset 1). This was also the case for proteins encoded in the sucrose/β-glucan genetic region, but only for the sucrose-related part (Fig. S3). Therefore, proteins encoded in the β-glucan part (GH16_3 and co-encoded TBDT) are regulated independently from the sucrose PUL. Protein abundance of the distal PdGH68_DIS_ and PdGH32_DIS_ was very low (%riBAQ <0.002 in fructan-grown cells, Fig. S4) compared to proteins encoded in the fructan and sucrose PUL. However, compared to whole-cell proteomes, PdGH68_DIS_ was the most abundant protein in the extracellular protein fraction of inulin-grown cells (Fig. S5, Supplementary Dataset 2).

Unexpectedly, abundance of proteins related to the utilization of α-glucans was increased in cells grown on bacterial levan (Fig. 2), but was much lower compared to α-glucan-grown cells (Fig. S6). *P. distincta* hardly grew on bacterial levan. The increased protein abundance of α-glucan-related proteins might indicate that *P. distincta* uses an α-glucan-type storage polysaccharide, like glycogen (42), when cultured with bacterial levan.

**Fig. 2.**
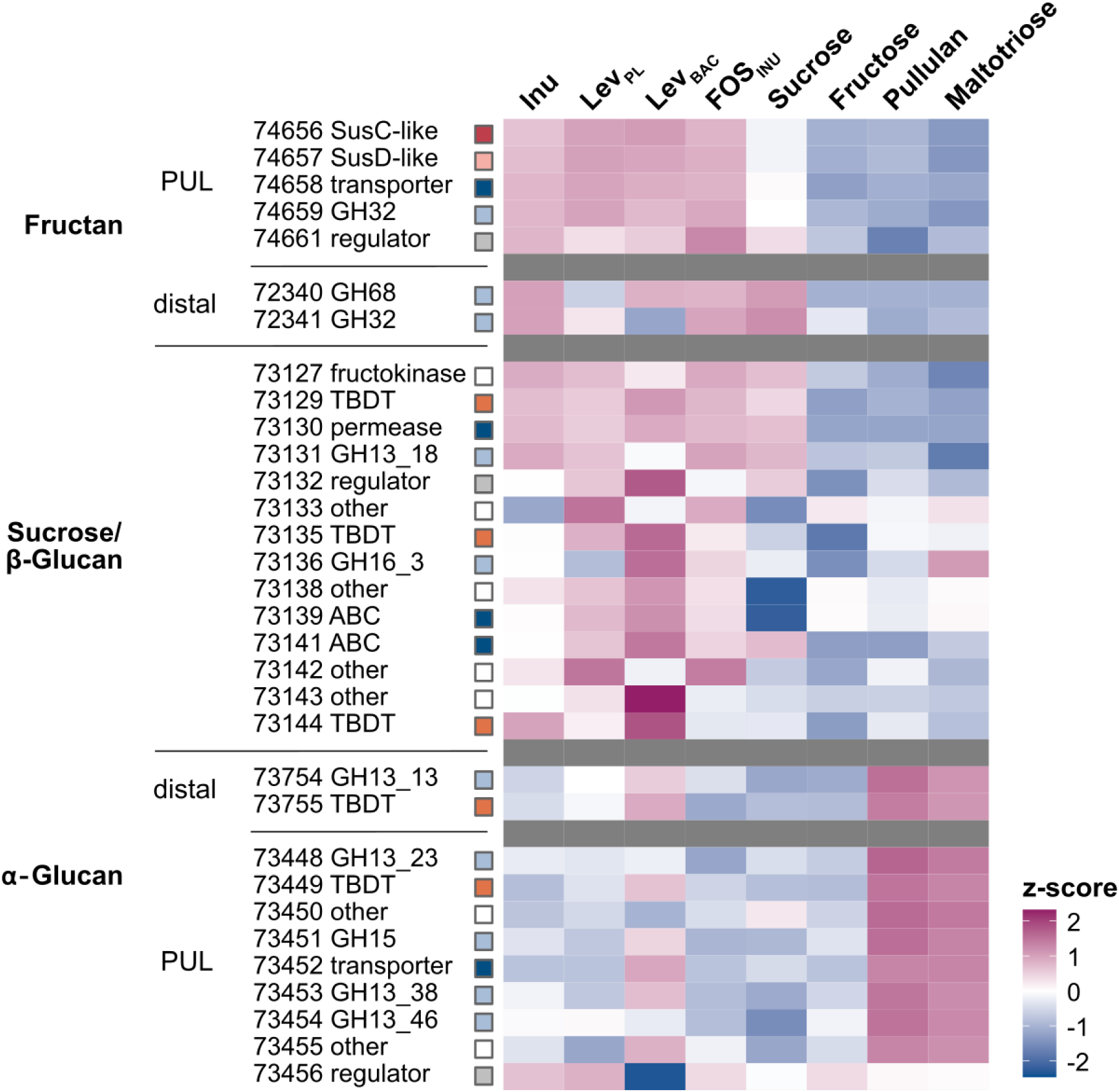
The abundance of fructan PUL-encoded proteins is increased in inulin-grown and levan-grown cells. Relative protein abundance of selected proteins of *P. distincta* grown with different fructose-containing substrates and α-glucans as control; indicated above the plot. Relative protein abundance values (%riBAQ, relative intensity-based absolute quantification, giving the relative protein abundance within the sample, n=3) were log-normalized and z-scored. Protein accessions were abbreviated, e.g., 74657 refers to EGI74657.1. Inu: Inulin, LevPL: Plant-derived levan, LevBAC: Bacterial levan, FOSINU: Fructo-oligosaccharides from inulin.

### The SusC/D-like complex in Pseudoalteromonas distincta is specific for inulin-like substrates

Structural and sequence comparisons showed that the *P. distincta* SusC/D-like proteins (PdSusC/D_PUL_) are indeed very similar to *Bacteroidota* fructan-specific homologs. For comparisons, we used proteins from human intestinal *Bacteroidota*, where fructan utilization is well-studied and 3D crystal structures of levan-specific proteins are available.

The *P. distincta* SusC/D-like proteins have a high sequence similarity to the *Bacteroidota* homologs known or suggested to bind/transport levan or inulin (PdSusD_PUL_: 53.5-61.5%, PdSusC_PUL_: 56.6-61.7%, see Supplementary Table S3, Fig. S7). Furthermore, the AF-predicted 3D structures of the *P. distincta* proteins superimpose very well with the corresponding levan-specific complex from *B. thetaiotaomicron* VPI-5482^T^ (PDB ID 6ZAZ, (63)) (Fig. 3). However, some levan-binding residues of *B. thetaiotaomicron* VPI-5482^T^ are missing in *P. distincta* (Fig. 3, Fig. S7) and in a large-scale phylogenetic analysis, PdSusD_PUL_ did not cluster with these selected *Bacteroidota* homologs (Fig. S8).

**Fig. 3.**
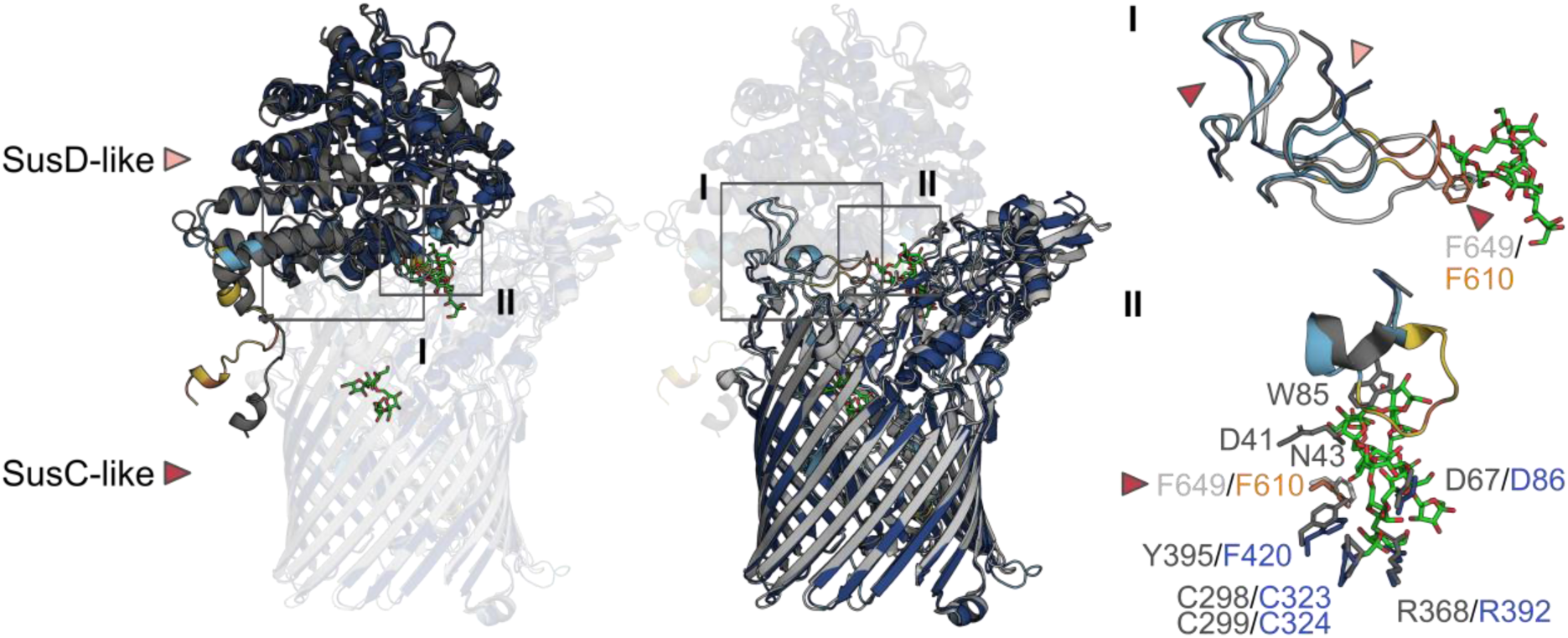
A SusC/D-like transport complex in *Pseudoalteromonas distincta*. Superposition of the AF-predicted 3D structures of PdSusC/DPUL, colored using the pLDDT confidence scores, with the corresponding levan-binding complex from *Bacteroides thetaiotaomicron* VPI-5482^T^ (PDB ID 6ZAZ, (63)) in gray. Boxed region “I” highlights ‘hinge’ loops responsible for the interaction between the glycan-binding SusD-like protein and SusC-like TBDT. Boxed region “II” shows residues of the SusD-like protein from *B. thetaiotaomicron* binding FOS from levan and their superposition with residues of the *P. distincta* PdSusDPUL (colored with confidence scores), if similar or identical, as a top view (see also Fig. S7). A levan-interacting residue of the SusC-like protein in *B. thetaiotaomicron* (64) – also shown in box “I” – is marked with an arrow head. pLDDT confidence scores: “very high” (pLDDT>90) dark blue, “high” (90> pLDDT>70) cyan, “low” (70> pLDDT>50) yellow and “very low” (pLDDT<50) orange.

We therefore investigated ligand binding by PdSusD_PUL_. Affinity gel electrophoresis (1% of glycan used) and surface plasmon resonance did not detect binding of PdSusD_PUL_ to levan or inulin (data not shown). Similarly, isothermal titration calorimetry (ITC) did not show binding to inulin, levan, sucrose or fructose at the concentrations used (data not shown, see Materials and Methods). However, PdSusD_PUL_ did show low affinity to FOS_INU_ (K_a_ ∼43 M^-1^, Fig. 4A, Fig. S9A). The interaction was enthalpically favorable and entropically unfavorable (Fig. S9B). Attempts to obtain a 3D crystal structure for PdSusD_PUL_ to gain further insight into ligand binding were not successful (see Supplementary Methods).

**Fig. 4.**
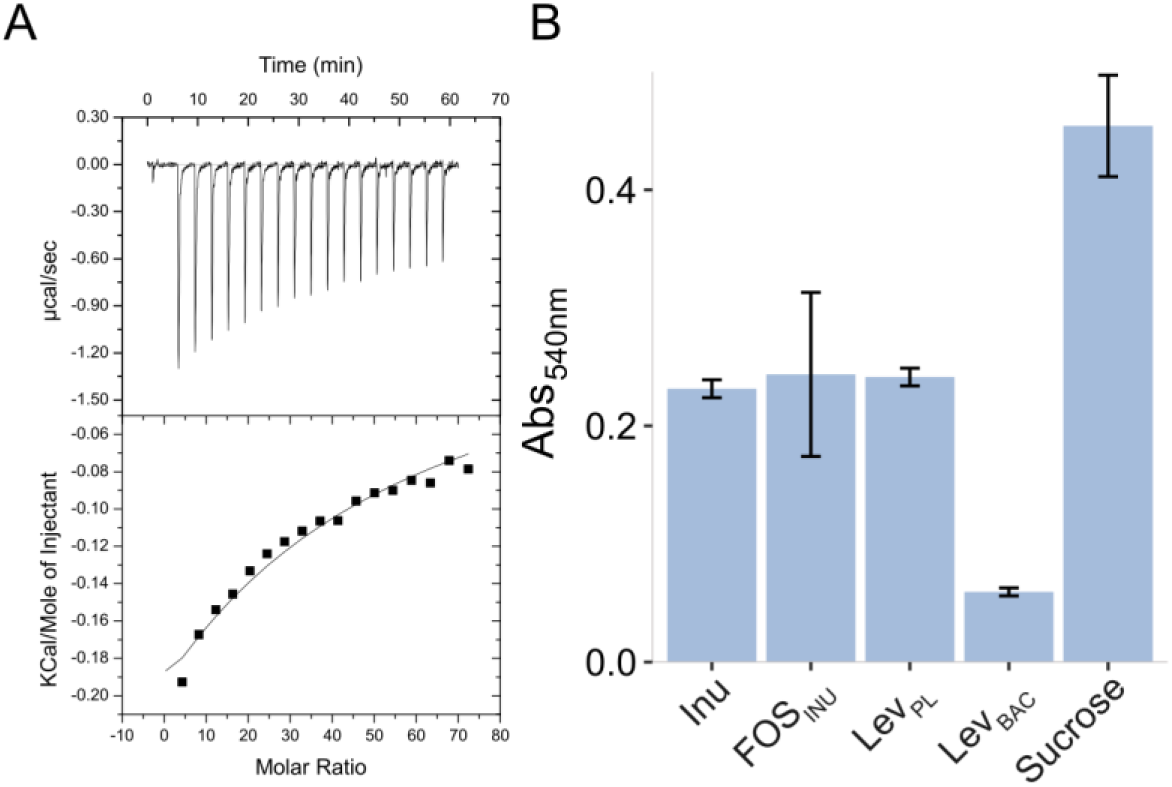
The PUL-encoded SusD-like protein binds FOSINU, while the PUL-encoded GH32 cleaves inulin, levan and sucrose. (**A**) Binding of PdSusDPUL to FOSINU. ITC raw titration data (upper panel) and integrated heat peaks (lower panel) as a function of the molar ratio of ligand per receptor. (**B**) Activity of PdGH32PUL on different fructans and sucrose as detected by RSA, shown as mean with standard deviation (n=3). Values were corrected against controls without CAZyme. As a reference for a non-active CAZyme see also Fig. S10A. Inu: Inulin, FOSINU: Fructo-oligosaccharides from inulin, LevPL: Plant-derived levan, LevBAC: Bacterial levan.

### The fructan PUL-encoded GH32 of Pseudoalteromonas distincta efficiently hydrolyzes inulin, but also plant-derived levan

The PUL-encoded CAZyme PdGH32_PUL_ hydrolyzed inulin, FOS_INU_, plant-derived levan and sucrose, but also showed low activity on bacterial levan, as determined by the reducing sugar assay (RSA) and the fluorophore-assisted carbohydrate electrophoresis (FACE) (Fig. 4B, Fig. S10A-D). FACE analysis suggested exo-activity of PdGH32_PUL_ as sucrose was still hydrolyzed (Fig. 4B, Fig. S10D). In line with this, AF predictions indicate a ‘funnel’-like binding pocket of PdGH32_PUL_ (Fig. S11). A special feature of the PUL-encoded PdGH32_PUL_ is an additional carbohydrate-binding module (CBM38), which protrudes between blade IV and V of the five-bladed β-propeller catalytic module (Fig. S12). Attempts to obtain 3D crystal structures of PdGH32_PUL_ were not successful (see Supplementary Methods).

In contrast to the PUL-encoded PdGH32_PUL_, the distal PdGH32_DIS_ of *P. distincta* is levan-specific (Fig. S13) and endo-active (Fig. S14). In agreement with this, the predicted catalytic groove of the distal PdGH32_DIS_ is larger compared to the PUL-encoded PdGH32_PUL_ (Fig. S11). Moreover, we detected high sucrose-hydrolyzing activity for PdGH68_DIS_ (Fig. S10A and D), but no production of higher molecular weight compounds.

### SusC/D-like proteins of Gammaproteobacteria are mostly associated to strains of marine origin and to fructan utilization

The phylogenetic analysis identified 13 additional gammaproteobacterial SusD-like proteins within a huge number of *Bacteroidota* homologs (1,005 in a total of 1,055 protein sequences; Fig. S8). Gammaproteobacterial sequences are of sediment/soil, aquatic and marine origin (Table S3) and are encoded in GH32-containing PULs (data not shown). To further investigate this, we searched the GEM and UGCMP databases for gammaproteobacterial SusD-like sequences. In a total of 4,681 representative *Gammaproteobacteria* genomes, 32 clusters (31 genomes) encoded SusD-like proteins, out of which 26 were of marine origin (Fig. 5). Most SusD-like protein-coding sequences occurred as SusC/D-like pairs (91%). Out of these SusC/D-like pairs, 83% co-occurred with GH32s, indicating fructan PULs (Fig. 5). The number of SusC/D-containing PULs might be even higher due to incompleteness of clusters (occasionally fragmented genomes).

In addition, 631 GH32s were identified in 574 gammaproteobacterial clusters (426 genomes) using this dataset. 4% of the identified GH32s are encoded in the above-mentioned SusC/D-containing PULs. 7% co-occurred with TBDTs instead, indicating classical *Gammaproteobacteria* PULs. 89% did not contain SusC/D-like proteins or TBDTs in their genomic neighborhood suggesting that many GH32 are not encoded in PULs.

**Fig. 5.**
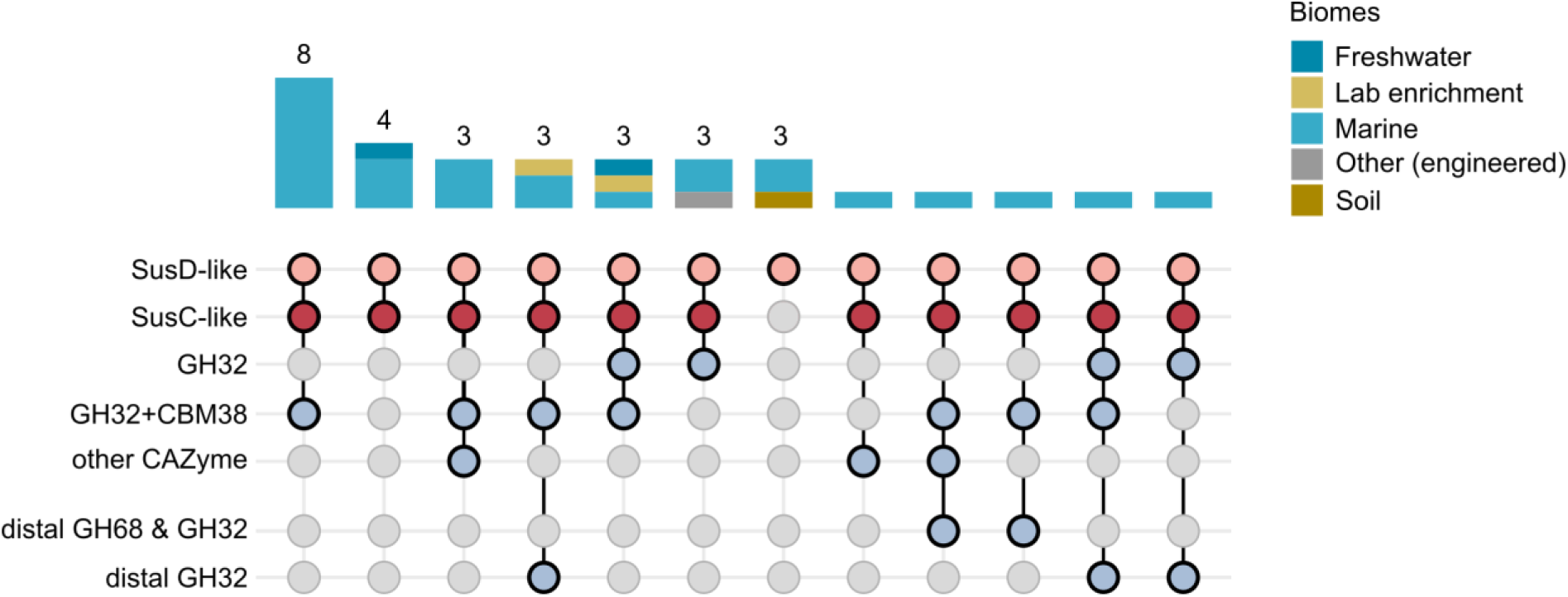
SusD-like sequences co-occur with SusC-like TBDTs and GH32s in *Gammaproteobacteria*. The GEM and UGCMP databases were searched for gammaproteobacterial SusD-like sequences (4,681 representative genomes). Shown is their genomic context as well as the presence of additional GH68/GH32 pairs or GH32s.

## Discussion

Here we show that marine *Gammaproteobacteria* such as *P. distincta* degrade fructans, which is enabled by a PUL encoding a GH32 and a SusC/D-like outer membrane transport complex. The presence of a glycan-binding SusD-like protein and a SusC-like TBDT in *P. distincta* is in contrast to the current state of research that this transport complex is restricted to the *Bacteroidota* phylum. The *P. distincta* proteins were thus probably acquired from a bacteroidotal ancestor.

Like SusD-like proteins in *Bacteroidota*, the fructan PUL-encoded SusD-like protein (PdSusD_PUL_) of *P. distincta* contains a lipoprotein signal peptide that directs export and membrane anchoring. Interestingly, in PdSusD_PUL_, the conserved cysteine is followed by acidic residues (Fig. S15), which signal lipoprotein export to the bacterial surface in *Bacteroidota* (65). In addition, the *P. distincta* SusC/D-like complex is structurally predicted to be highly similar to the levan-binding SusC/D-like proteins from *B. thetaiotaomicron* VPI-5482^T^ (3, 63). This similarity includes specific loops at the top of the SusC β-barrel and close to the fructan binding site of the SusD-like protein that mediate interaction between the two proteins (Fig. 3). In comparison, the gammaproteobacterial SusC-like proteins identified in this study lack the N-terminal carboxypeptidase D regulatory-like domain. The exact function of the carboxypeptidase D regulatory-like domain preceding the plug domain is unknown, but many SusC-like TBDTs of *B. thetaiotaomicron* VPI-5482^T^ possess it (66). This includes the levan-specific protein (BT_1763) where it was shown that the deletion of the carboxypeptidase D regulatory-like domain causes a growth defect (63). It was speculated that it might mediate specificity for the various TonB orthologs. With one exception (EGI71291), all the other TBDTs of *P. distincta* also lack the carboxypeptidase D regulatory-like domain, suggesting a functional complex despite the missing domain. Horizontal transfers of genes or operons related to polysaccharide use have been described before, from marine *Bacteroidota* to *Bacteroidota* of the human intestine (67, 68), but also from a marine flavobacterium (*Bacteroidota*) to *Proteobacteria*, including *Gammaproteobacteria* (69). However, in the latter case, the transfer of *Bacteroidota* alginolytic operons to *Gammaproteobacteria* did not include the SusC/D-like sequences. It is therefore intriguing how the fructan-related complex was able to establish in *Gammaproteobacteria*, while SusC/Ds with other specificities did not – or not yet.

While we could not show binding of the *P. distincta* PdSusD_PUL_ to levan and inulin, we confirmed its binding to FOS_INU_. Together with the much better growth of *P. distincta* on inulin, this suggests that the SusC/D-like complex transports inulin-like fructans. In human gastrointestinal *Bacteroidota*, it has been shown that the specificity towards levan or inulin is mediated by SusC/D-like proteins and by the subcellular localization of GH32s (11). At the same time, the affinity of PdSusD_PUL_ to FOS_INU_ was much lower (∼K_a_ 43 M^-1^) compared to the affinity of the inulin-binding SusD-like protein of *B. thetaiotaomicron* strain Bt-8736 (the K_d_ was reported as ∼0.5 mM, which corresponds to K_a_ 2 x 10^3^ M^-1^) (11). It is possible that higher affinity binding requires the functional complex with the SusC-like TBDT, as already speculated for a non-binding SusD-like protein encoded in a xylo-oligosaccharide PUL (70).

In addition to PdSusD_PUL_, the fructan PUL-encoded CAZyme PdGH32_PUL_ contains a lipoprotein signal peptide. However, the conserved cysteine in PdGH32_PUL_ is not followed by acidic residues (Fig. S15) indicating that the CAZyme is not exposed to the surface. Together with its high protein abundance in the outer membrane and outer membrane vesicles (Fig. S5, Supplementary Data Set 2), this suggests that PdGH32_PUL_ is anchored to the outer membrane, but orientated towards the periplasm. Therefore, despite the PdGH32_PUL_ activity on inulin and levan, known for exo-acting GH32s (8), only inulin would enter - thanks to the SusC/D_PUL_ specificity - and thus only inulin would be digested by PdGH32_PUL_ (Fig. 6). Similarly, inulin-degrading *Bacteroidota* from the human intestine lack extracellular GH32s or extracellular CAZymes are not necessary for growth (11, 71). However, in *P. distincta*, growth on levan from plants may be enabled by the extracellular PdGH32_DIS_ (Fig. S5, Supplementary Data Set 2), which is not encoded in a PUL. Bacterial levan has a high molecular weight (>1000 kDa (72)) and is branched (73) compared to plant-derived levan (∼12.5 kDa), which could explain differences seen in growth on levans.

An additional GH68/GH32 pair in *P. distincta* and other *Gammaproteobacteria* of marine origin (Fig. 5) (74) indicates fructan production (GH68) (6) and recycling/remodeling (GH32). We demonstrated sucrose hydrolysis by PdGH68_DIS_, but could not confirm the production of higher molecular weight FOS or fructans. However, using the AF predicted structure of PdGH68_DIS_ and DALI (75) for structural comparisons of proteins, the GH68 of *Microbacterium saccharophilum* K-1 (formerly *Arthrobacter*, z-score 66.2, identity 77%, PDB ID 3VSS (76)) and the GH68 of *Gluconacetobacter diazotrophicus* SRT4 (formerly *Acetobacter diazotrophicus*, z-score 61.3, identity 56%, PDB ID 1W18 (77)) were identified as closest homologs, both with transfructosylation activities (78, 79). The production of fructans and fructose-containing EPS by marine bacteria has been reported before (80, 81). Although the transfructosylation activity of PdGH68_DIS_ has not yet been demonstrated, this indicates fructose-based EPS as one source for marine fructans.

**Fig. 6.**
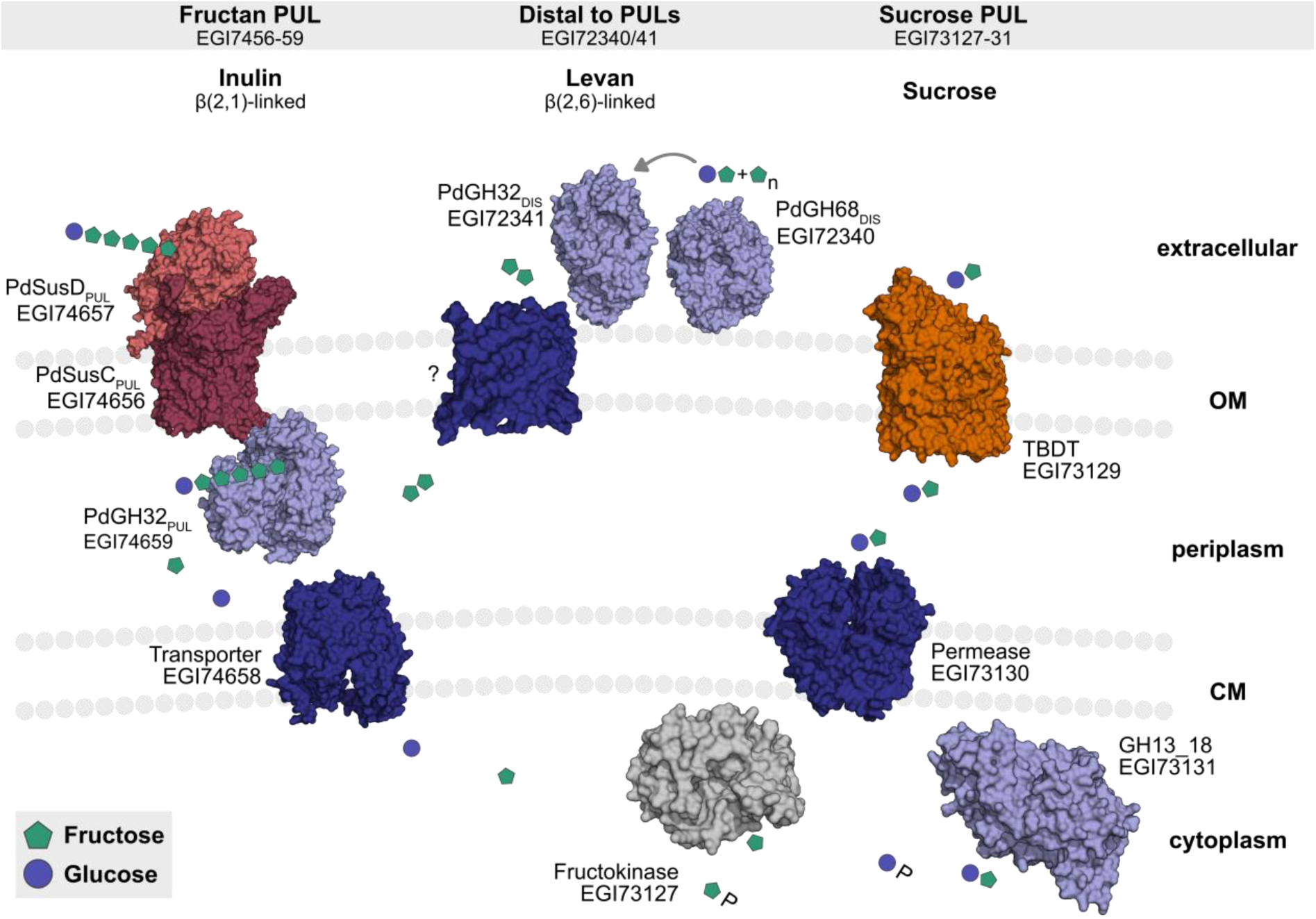
Proteins that are encoded in different genomic regions mediate the utilization of inulin, levan or sucrose, but could be linked during downstream processing. Our model for the utilization of fructose-containing substrates in *Pseudoalteromonas distincta* is presented. Left: Fructan PUL-encoded proteins enable inulin use. Presumably, the small-sized inulin is shuttled directly into the periplasm by the SusC/D-like transport complex, where it is degraded by the exo-active CAZyme PdGH32PUL, as previously described for human gastrointestinal *Bacteroidota* (11). Top center: A distal PdGH68DIS could produce levan as EPS, this levan could be degraded/remodelled by the endo-levanase PdGH32DIS. Fragments released from PdGH32DIS might be further degraded in the periplasm by the fructan PUL-encoded PdGH32PUL, which accepts both linkages. The transporter importing these oligosaccharides is unknown. Right: Proteins encoded in a sucrose PUL may metabolize fructose and sucrose, released upon degradation of inulin and levan. In inulin and levan, fructosyl units are β(2,1)-or β(2,6)-linked to sucrose, respectively.

Fructan utilization may be more widespread than suggested by the frequency of fructan PULs detected in our analysis and in bacterial metagenome-assembled genomes from phytoplankton blooms (16), because GH32s often occur without a PUL context. However, GH32-containing PULs may have the advantage that the expression is fine-tuned by regulatory elements. Overall, it remains speculative, if the only benefit of acquiring a fructan PUL from a putative *Bacteroidota* ancestor was an extended substrate repertoire or if the SusC/D-like tandem came with advantageous aspects. For instance, for *Porphyromonas gingivalis*, a pathogen *Bacteroidota* in the human periodontitis, it was suggested that the peptide-binding RagB-like protein (corresponding to SusD-like protein) mediates specificity (61). In addition, a SusD-like protein might be beneficial, when endo-active extracellular inulinases and CBM-containing proteins are missing, as in *P. distincta*.

The discovery of SusC/D-like proteins in fructan PULs of marine *Gammaproteobacteria* indicates an exchange of protein functions between phyla to support glycan utilization. Our findings show that fructose-containing substrates contribute to the marine glycan universe targeted by specialized *Gammaproteobacteria* such as *P. distincta*.

## Supporting information

Supporting Informaton

## Data Availability

The proteomic data were deposited on the PRIDE archive. Data will be released upon acceptance of the manuscript in a peer-reviewed journal.

## Acknowledgements

The authors thank Jana Matulla, Theresa Dutschei, Julia-Maria Seeßelberg and Marcella Consolo for support in the lab, Tjorven Hinzke and Finn Lillelund Aachmann for helpful comments as well as Sebastian Grund for LC-MS/MS measurements. We are grateful for having access to the CristalO platform (FR2424, Station Biologique de Roscoff), which is part of the core facility networks Biogenouest and EMBRC-France, and to the P11 beamline at DESY (Hamburg, Germany). We thank the staff for their support on site. MKZ thanks the DFG (SCHW 595/10-3), which enabled a research stay in Roscoff.

## Author contributions

MKZ and TS designed and directed the study. MKZ, AB, VS, MT, NW, FU, EFB, PS, LS, LC, AJ, AS, FH, DJ, MJ and MC performed research. MKZ, DBa, EFB, TB, MH and MC analyzed the data. MD, UTB, DBe, JHH, MC and TS provided resources. MKZ wrote the paper, which was edited and approved by all authors.

## References

1. Anderson KL, Salyers AA. Biochemical evidence that starch breakdown by *Bacteroides thetaiotaomicron* involves outer membrane starch-binding sites and periplasmic starch-degrading enzymes. J Bacteriol. 1989;171(6):3192–8; doi:10.1128/jb.171.6.3192-3198.1989.

2. Bjursell MK, Martens EC, Gordon JI. Functional genomic and metabolic studies of the adaptations of a prominent adult human gut symbiont, *Bacteroides thetaiotaomicron*, to the suckling period. J Biol Chem. 2006;281(47):36269–79; doi:10.1074/jbc.M606509200.

3. Glenwright AJ, Pothula KR, Bhamidimarri SP, Chorev DS, Baslé A, Firbank SJ, et al. Structural basis for nutrient acquisition by dominant members of the human gut microbiota. Nature. 2017;541(7637):407-11; doi:10.1038/nature20828.

4. Bolam DN, van den Berg B. TonB-dependent transport by the gut microbiota: novel aspects of an old problem. Curr Opin Struct Biol. 2018;51:35–43; doi:10.1016/j.sbi.2018.03.001.

5. Lammens W, Le Roy K, Schroeven L, Van Laere A, Rabijns A, Van den Ende W. Structural insights into glycoside hydrolase family 32 and 68 enzymes: functional implications. J Exp Bot. 2009;60(3):727–40; doi:10.1093/jxb/ern333.

6. Versluys M, Kirtel O, Toksoy Öner E, Van den Ende W. The fructan syndrome: Evolutionary aspects and common themes among plants and microbes. Plant Cell Environ. 2018;41(1):16–38; doi:10.1111/pce.13070.

7. Zhang WL, Xu W, Ni DW, Dai QY, Guang CE, Zhang T, et al. An overview of levan-degrading enzyme from microbes. Appl Microbiol Biotechnol. 2019;103(19):7891–902; doi:10.1007/s00253-019-10037-4.

8. Sonnenburg ED, Zheng HJ, Joglekar P, Higginbottom SK, Firbank SJ, Bolam DN, et al. Specificity of polysaccharide use in intestinal *Bacteroides* species determines diet-induced microbiota alterations. Cell. 2010;141(7):1241–U256; doi:10.1016/j.cell.2010.05.005.

9. Ernits K, Eek P, Lukk T, Visnapuu T, Alarnäe T. First crystal structure of an endo-levanase - the BT1760 from a human gut commensal *Bacteroides thetaiotaomicron*. Sci Rep. 2019;9; doi:10.1038/s41598-019-44785-0.

10. Mardo K, Visnapuu T, Vija H, Aasamets A, Viigand K, Alamae T. A Highly Active Endo-Levanase BT1760 of a Dominant Mammalian Gut Commensal *Bacteroides thetaiotaomicron* Cleaves Not Only Various Bacterial Levans, but Also Levan of Timothy Grass. Plos One. 2017;12(1):e0169989; doi:10.1371/journal.pone.0169989.

11. Joglekar P, Sonnenburg ED, Higginbottom SK, Earle KA, Morland C, Shapiro-Ward S, et al. Genetic Variation of the SusC/SusD Homologs from a Polysaccharide Utilization Locus Underlies Divergent Fructan Specificities and Functional Adaptation in *Bacteroides thetaiotaomicron* Strains. mSphere. 2018;3(3); doi:10.1128/mSphereDirect.00185-18.

12. Li AX, Guo LZ, Lu WD. Alkaline inulinase production by a newly isolated bacterium Marinimicrobium sp LS-A18 and inulin hydrolysis by the enzyme. World J Microbiol Biotechnol. 2012;28(1):81–9; doi:10.1007/s11274-011-0794-3.

13. Gao L, Chi Z, Sheng J, Wang L, Li J, Gong F. Inulinase-producing marine yeasts: evaluation of their diversity and inulin hydrolysis by their crude enzymes. Microb Ecol. 2007;54(4):722–9; doi:10.1007/s00248-007-9231-4.

14. 14. Abd El Aty AA, Wehaidy HR, Mostafa FA. Optimization of inulinase production from low cost substrates using Plackett-Burman and Taguchi methods. Carbohydr Polym. 2014;102:261–8; doi:10.1016/j.carbpol.2013.11.007.

15. Liebl W, Brem D, Gotschlich A. Analysis of the gene for beta-fructosidase (invertase, inulinase) of the hyperthermophilic bacterium *Thermotoga maritima*, and characterisation of the enzyme expressed in *Escherichia coli*. Appl Microbiol Biotechnol. 1998;50(1):55–64; doi:10.1007/s002530051256.

16. Wang FQ, Bartosik D, Sidhu C, Siebers R, Lu DC, Trautwein-Schult A, et al. Particle-attached bacteria act as gatekeepers in the decomposition of complex phytoplankton polysaccharides. Microbiome. 2024;12(32); doi:10.1186/s40168-024-01757-5.

17. Sidhu C, Kirstein IV, Meunier CL, Rick J, Fofonova V, Wiltshire KH, et al. Dissolved storage glycans shaped the community composition of abundant bacterioplankton clades during a North Sea spring phytoplankton bloom. Microbiome. 2023;11(1):77; doi:10.1186/s40168-023-01517-x.

18. Alberto F, Jordi E, Henrissat B, Czjzek M. Crystal structure of inactivated invertase in complex with the trisaccharide substrate raffinose. Biochem J. 2006;395:457–62; doi:10.1042/Bj20051936.

19. Schut F, de Vries EJ, Gottschal JC, Robertson BR, Harder W, Prins RA, et al. Isolation of typical marine bacteria by dilution culture: growth, maintenance, and characteristics of isolates under laboratory conditions. Appl Environ Microbiol. 1993;59(7):2150–60; doi:10.1128/aem.59.7.2150-2160.1993.

20. Dürwald A, Zühlke MK, Schlüter R, Gebbe R, Bartosik D, Unfried F, et al. Reaching out in anticipation: bacterial membrane extensions represent a permanent investment in polysaccharide sensing and utilization. Environ Microbiol. 2021;23(6):3149–63; doi:10.1111/1462-2920.15537.

21. Frias A, Manresa A, de Oliveira E, López-Iglesias C, Mercade E. Membrane vesicles: a common feature in the extracellular matter of cold-adapted Antarctic bacteria. Microb Ecol. 2010;59(3):476–86; doi:10.1007/s00248-009-9622-9.

22. McBroom AJ, Johnson AP, Vemulapalli S, Kuehn MJ. Outer membrane vesicle production by *Escherichia coli* is independent of membrane instability. J Bacteriol. 2006;188(15):5385–92; doi:10.1128/Jb.00498-06.

23. Bonn F, Otto A. Protein Enrichment from Highly Dilute Samples with StrataClean. Methods Mol Biol. 2018;1841:11–8; doi:10.1007/978-1-4939-8695-8_2.

24. Bonn F, Bartel J, Büttner K, Hecker M, Otto A, Becher D. Picking vanished proteins from the void: how to collect and ship/share extremely dilute proteins in a reproducible and highly efficient manner. Anal Chem. 2014;86(15):7421–7; doi:10.1021/ac501189j.

25. Cox J, Mann M. MaxQuant enables high peptide identification rates, individualized p.p.b.-range mass accuracies and proteome-wide protein quantification. Nat Biotechnol. 2008;26(12):1367–72; doi:10.1038/nbt.1511.

26. Tyanova S, Temu T, Sinitcyn P, Carlson A, Hein MY, Geiger T, et al. The Perseus computational platform for comprehensive analysis of (prote)omics data. Nat Methods. 2016;13(9):731; doi:10.1038/nmeth.3901.

27. Tyanova S, Cox J. Perseus: a bioinformatics platform for integrative analysis of proteomics data in cancer research. Methods Mol Biol. 2018;1711:133–48; doi:10.1007/978-1-4939-7493-1_7.

28. Vizcaíno JA, Csordas A, del-Toro N, Dianes JA, Griss J, Lavidas I, et al. 2016 update of the PRIDE database and its related tools. Nucleic Acids Res. 2016;44(D1):D447–56; doi:10.1093/nar/gkv1145.

29. Perez-Riverol Y, Bai J, Bandla C, García-Seisdedos D, Hewapathirana S, Kamatchinathan S, et al. The PRIDE database resources in 2022: a hub for mass spectrometry-based proteomics evidences. Nucleic Acids Res. 2022;50(D1):D543–D52; doi:10.1093/nar/gkab1038.

30. Yu NY, Wagner JR, Laird MR, Melli G, Rey S, Lo R, et al. PSORTb 3.0: improved protein subcellular localization prediction with refined localization subcategories and predictive capabilities for all prokaryotes. Bioinformatics. 2010;26(13):1608–15; doi:10.1093/bioinformatics/btq249.

31. Moreno J, Nielsen H, Winther O, Teufel F. Predicting the subcellular location of prokaryotic proteins with DeepLocPro. Bioinformatics. 2024;40(12); doi:10.1093/bioinformatics/btae677.

32. Teufel F, Almagro Armenteros JJ, Johansen AR, Gíslason MH, Pihl SI, Tsirigos KD, et al. SignalP 6.0 predicts all five types of signal peptides using protein language models. Nat Biotechnol. 2022;40(7):1023–5; doi:10.1038/s41587-021-01156-3.

33. Savojardo C, Martelli PL, Fariselli P, Casadio R. DeepSig: deep learning improves signal peptide detection in proteins. Bioinformatics. 2018;34(10):1690–6; doi:10.1093/bioinformatics/btx818.

34. Gibson DG, Young L, Chuang RY, Venter JC, Hutchison CA, 3rd, Smith HO. Enzymatic assembly of DNA molecules up to several hundred kilobases. Nat Methods. 2009;6(5):343–5; doi:10.1038/nmeth.1318.

35. Studier FW. Protein production by auto-induction in high-density shaking cultures. Protein Expr Purif. 2005;41(1):207–34; doi:10.1016/j.pep.2005.01.016.

36. Hehemann JH, Smyth L, Yadav A, Vocadlo DJ, Boraston AB. Analysis of Keystone Enzyme in Agar Hydrolysis Provides Insight into the Degradation of a Polysaccharide from Red Seaweeds. J Biol Chem. 2012;287(17):13985–95; doi:10.1074/jbc.M112.345645.

37. Wilkins MR, Gasteiger E, Bairoch A, Sanchez JC, Williams KL, Appel RD, et al. Protein identification and analysis tools in the ExPASy server. Methods Mol Biol. 1999;112:531–52; doi:10.1385/1-59259-584-7:531.

38. Wiseman T, Williston S, Brandts JF, Lin LN. Rapid Measurement of Binding Constants and Heats of Binding Using a New Titration Calorimeter. Anal Biochem. 1989;179(1):131–7; doi:10.1016/0003-2697(89)90213-3.

39. Varadi M, Anyango S, Deshpande M, Nair S, Natassia C, Yordanova G, et al. AlphaFold Protein Structure Database: massively expanding the structural coverage of protein-sequence space with high-accuracy models. Nucleic Acids Res. 2022;50(D1):D439–D44; doi:10.1093/nar/gkab1061.

40. Consortium TU. UniProt: the universal protein knowledgebase in 2021. Nucleic Acids Res. 2021;49(D1):D480–D9; doi:10.1093/nar/gkaa1100.

41. Altschul SF, Gish W, Miller W, Myers EW, Lipman DJ. Basic Local Alignment Search Tool. J Mol Biol. 1990;215(3):403–10; doi:DOI 10.1006/jmbi.1990.9999.

42. Barbeyron T, Thomas F, Barbe V, Teeling H, Schenowitz C, Dossat C, et al. Habitat and taxon as driving forces of carbohydrate catabolism in marine heterotrophic bacteria: example of the model algae-associated bacterium *Zobellia galactanivorans* Dsij^T^. Environ Microbiol. 2016;18(12):4610–27; doi:10.1111/1462-2920.13584.

43. Ficko-Blean E, Préchoux A, Thomas F, Rochat T, Larocque R, Zhu YT, et al. Carrageenan catabolism is encoded by a complex regulon in marine heterotrophic bacteria. Nat Commun. 2017;8; doi:10.1038/s41467-017-01832-6.

44. Katoh K, Misawa K, Kuma K, Miyata T. MAFFT: a novel method for rapid multiple sequence alignment based on fast Fourier transform. Nucleic Acids Res. 2002;30(14):3059–66; doi:10.1093/nar/gkf436.

45. Clamp M, Cuff J, Searle SM, Barton GJ. The Jalview Java alignment editor. Bioinformatics. 2004;20(3):426–7; doi:10.1093/bioinformatics/btg430.

46. Felsenstein J. Evolutionary Trees from DNA-Sequences - a Maximum-Likelihood Approach. J Mol Evol. 1981;17(6):368–76; doi:10.1007/Bf01734359.

47. Stamatakis A. RAxML version 8: a tool for phylogenetic analysis and post-analysis of large phylogenies. Bioinformatics. 2014;30(9):1312–3; doi:10.1093/bioinformatics/btu033.

48. Trifinopoulos J, Nguyen LT, von Haeseler A, Minh BQ. W-IQ-TREE: a fast online phylogenetic tool for maximum likelihood analysis. Nucleic Acids Res. 2016;44(W1):W232–W5; doi:10.1093/nar/gkw256.

49. Le SQ, Gascuel O. An improved general amino acid replacement matrix. Mol Biol Evol. 2008;25(7):1307–20; doi:10.1093/molbev/msn067.

50. Felsenstein J. Confidence-Limits on Phylogenies - an Approach Using the Bootstrap. Evolution. 1985;39(4):783–91; doi:10.2307/2408678.

51. Letunic I, Bork P. Interactive Tree Of Life (iTOL) v5: an online tool for phylogenetic tree display and annotation. Nucleic Acids Res. 2021;49(W1):W293–W6; doi:10.1093/nar/gkab301.

52. Nishimura Y, Yoshizawa S. The OceanDNA MAG catalog contains over 50,000 prokaryotic genomes originated from various marine environments. Sci Data. 2022;9(1):305; doi:10.1038/s41597-022-01392-5.

53. Nayfach S, Roux S, Seshadri R, Udwary D, Varghese N, Schulz F, et al. A genomic catalog of Earth’s microbiomes. Nat Biotechnol. 2021;39(4):499–509; doi:10.1038/s41587-020-0718-6.

54. Olm MR, Brown CT, Brooks B, Banfield JF. dRep: a tool for fast and accurate genomic comparisons that enables improved genome recovery from metagenomes through de-replication. ISME J. 2017;11(12):2864–8; doi:10.1038/ismej.2017.126.

55. Seemann T. Prokka: rapid prokaryotic genome annotation. Bioinformatics. 2014;30(14):2068–9; doi:10.1093/bioinformatics/btu153.

56. Eddy SR. Accelerated Profile HMM Searches. PLoS Comp Biol. 2011;7(10):e1002195; doi:10.1371/journal.pcbi.1002195.

57. Mistry J, Chuguransky S, Williams L, Qureshi M, Salazar GA, Sonnhammer ELL, et al. Pfam: The protein families database in 2021. Nucleic Acids Res. 2021;49(D1):D412–D9; doi:10.1093/nar/gkaa913.

58. Haft DH, Selengut JD, White O. The TIGRFAMs database of protein families. Nucleic Acids Res. 2003;31(1):371–3; doi:10.1093/nar/gkg128.

59. Zheng J, Hu B, Zhang X, Ge Q, Yan Y, Akresi J, et al. dbCAN-seq update: CAZyme gene clusters and substrates in microbiomes. Nucleic Acids Res. 2023;51(D1):D557–D63; doi:10.1093/nar/gkac1068.

60. Lex A, Gehlenborg N, Strobelt H, Vuillemot R, Pfister H. UpSet: Visualization of Intersecting Sets. Ieee T Vis Comput Gr. 2014;20(12):1983–92; doi:10.1109/Tvcg.2014.2346248.

61. Madej M, White JBR, Nowakowska Z, Rawson S, Scavenius C, Enghild JJ, et al. Structural and functional insights into oligopeptide acquisition by the RagAB transporter from *Porphyromonas gingivalis*. Nat Microbiol. 2020;5(8):1016–25; doi:10.1038/s41564-020-0716-y.

62. Viborg AH, Terrapon N, Lombard V, Michel G, Czjzek M, Henrissat B, et al. A subfamily roadmap of the evolutionarily diverse glycoside hydrolase family 16 (GH16). J Biol Chem. 2019;294(44):15973–86; doi:10.1074/jbc.RA119.010619.

63. Gray DA, White JBR, Oluwole AO, Rath P, Glenwright AJ, Mazur A, et al. Insights into SusCD-mediated glycan import by a prominent gut symbiont. Nat Commun. 2021;12(1):44; doi:10.1038/s41467-020-20285-y.

64. White JBR, Silale A, Feasey M, Heunis T, Zhu YL, Zheng H, et al. Outer membrane utilisomes mediate glycan uptake in gut *Bacteroidetes*. Nature. 2023; doi:10.1038/s41586-023-06146-w.

65. Lauber F, Cornelis GR, Renzi F. Identification of a New Lipoprotein Export Signal in Gram-Negative Bacteria. Mbio. 2016;7(5); doi:10.1128/mBio.01232-16.

66. Pollet RM, Martin LM, Koropatkin NM. TonB-dependent transporters in the *Bacteroidetes*: Unique domain structures and potential functions. Mol Microbiol. 2021;115(3):490–501; doi:10.1111/mmi.14683.

67. Hehemann JH, Correc G, Barbeyron T, Helbert W, Czjzek M, Michel G. Transfer of carbohydrate-active enzymes from marine bacteria to Japanese gut microbiota. Nature. 2010;464(7290):908-12; doi:10.1038/nature08937.

68. Pudlo NA, Pereira GV, Parnami J, Cid M, Markert S, Tingley JP, et al. Diverse events have transferred genes for edible seaweed digestion from marine to human gut bacteria. Cell Host Microbe. 2022;30(3):314-+; doi:10.1016/j.chom.2022.02.001.

69. Thomas F, Barbeyron T, Tonon T, Génicot S, Czjzek M, Michel G. Characterization of the first alginolytic operons in a marine bacterium: from their emergence in marine *Flavobacteriia* to their independent transfers to marine Proteobacteria and human gut *Bacteroides*. Environ Microbiol. 2012;14(9):2379–94; doi:10.1111/j.1462-2920.2012.02751.x.

70. Tauzin AS, Wang Z, Cioci G, Li XQ, Labourel A, Machado B, et al. Structural and Biochemical Characterization of a Nonbinding SusD-Like Protein Involved in Xylooligosaccharide Utilization by an Uncultured Human Gut *Bacteroides* Strain. Msphere. 2022;7(5); doi:10.1128/msphere.00244-22.

71. Rakoff-Nahoum S, Foster KR, Comstock LE. The evolution of cooperation within the gut microbiota. Nature. 2016;533(7602):255-9; doi:10.1038/nature17626.

72. Keith J, Wiley B, Ball D, Arcidiacono S, Zorfass D, Mayer J, et al. Continuous culture system for production of biopolymer levan using *Erwinia herbicola*. Biotechnol Bioeng. 1991;38(5):557–60; doi:10.1002/bit.260380515.

73. Vijn I, Smeekens S. Fructan: More than a reserve carbohydrate? Plant Physiol. 1999;120(2):351–9; doi:DOI 10.1104/pp.120.2.351.

74. Gobet A, Barbeyron T, Matard-Mann M, Magdelenat G, Vallenet D, Duchaud E, et al. Evolutionary Evidence of Algal Polysaccharide Degradation Acquisition by *Pseudoalteromonas carrageenovora* 9^T^ to Adapt to Macroalgal Niches. Front Microbiol. 2018;9; doi:10.3389/fmicb.2018.02740.

75. Käll L, Krogh A, Sonnhammer EL. A combined transmembrane topology and signal peptide prediction method. J Mol Biol. 2004;338(5):1027–36; doi:10.1016/j.jmb.2004.03.016.

76. Tonozuka T, Tamaki A, Yokoi G, Miyazaki T, Ichikawa M, Nishikawa A, et al. Crystal structure of a lactosucrose-producing enzyme, *Arthrobacter* sp. K-1 β-fructofuranosidase. Enzyme Microb Technol. 2012;51(6-7):359–65; doi:10.1016/j.enzmictec.2012.08.004.

77. Martínez-Fleites C, Ortíz-Lombardía M, Pons T, Tarbouriech N, Taylor EJ, Arrieta JG, et al. Crystal structure of levansucrase from the Gram-negative bacterium *Gluconacetobacter diazotrophicus*. Biochem J. 2005;390:19–27; doi:10.1042/Bj20050324.

78. Hernandez L, Arrieta J, Menendez C, Vazquez R, Coego A, Suarez V, et al. Isolation and Enzymatic-Properties of Levansucrase Secreted by Acetobacter-Diazotrophicus Srt4, a Bacterium Associated with Sugar-Cane. Biochem J. 1995;309:113–8; doi:DOI 10.1042/bj3090113.

79. Ike M, Chen M, Danzl E, Sei K, Fujita M. Biodegradation of a variety of bisphenols under aerobic and anaerobic conditions. Water Sci Technol. 2006;53(6):153–9;

80. Okutani K. Structural Investigation of the Fructan from Marine Bacterium Nam-1. Bull Jap Soc Sci Fish. 1982;48(11):1621–5; doi:10.2331/suisan.48.1621.

81. Daly G, Decorosi F, Viti C, Adessi A. Shaping the phycosphere: Analysis of the EPS in diatom-bacterial co-cultures. J Phycol. 2023;59(4):791–7; doi:10.1111/jpy.13361.

