## Supporting Informaton for "Marine *Gammaproteobacteria* use SusC/D-like proteins in fructan utilization"

[*] corresponding authors

**The Supporting Information file contains**

Supplementary Methods

Supplementary Figures S1-S15

Supplementary Table S1-S3

Supplementary Data Sets 1 and 2 are provided as separate files

Supplementary Data Set 1: Whole-cell proteomes of *Pseudoalteromonas distincta*

Supplementary Data Set 2: Subcellular protein fractions of *Pseudoalteromonas distincta*

### Supplementary Methods

*Crystallization, data collection and data processing trials of PdSusD$_{PUL}$ and PdGH32$_{PUL}$*

Post-SEC purification, both proteins were used for initial crystallization attempts. Crystallization was performed in two drop 96-well crystallization plates in sitting drop format using 20 different commercial crystallization screens (more than 1800 conditions). The plates were setup with plausible substrates of the proteins, i.e. for PdGH32$_{PUL}$, levan oligosaccharides and bacterial/plant derived levan. PdSusD$_{PUL}$ with inulin and FOS$_{INU}$. Plates were incubated at 16° C. Start of the crystal formation was observed very late (~6 to 12 months). Diffraction quality crystals of PdSusD$_{PUL}$ were obtained in very few crystallization conditions (Structure screen F11 well and MCSG-2 screen H5 well). On the other hand, crystals of PdGH32$_{PUL}$ were obtained in many crystallization conditions. Cryo-protection in crystallization solution supplemented with 20% glycerol was done prior to flash cooling in a liquid nitrogen at 100 K. Diffraction data was collected on DESY beamline P11. A total of 3600 images were collected (exposure: 0.1s; oscillation: 0.1º). Repeated attempts were made with measured reflections intensities to index, integrate and scale using different image processing tools and pipeline available at the beamline facility and in-house facility, but we couldn't solve the structure for both proteins.

### Supplementary Tables and Figures

#### Table S1. Primer list.

| Primer | Sequence |
|---|---|
| Fw-pET28a | AAGCTTGCGGCCGCACTCGAG |
| Rev-pET28a | CATATGGCTGCCGCGCGGCAC |
| Fw-PdSusD$_{PUL}$ | TGCCGCGCGGCAGCCATATGTGTGGTGATTCTGATCTTGA |
| Rev-PdSusD$_{PUL}$ | AGTGCGGCCGCAAGCTTTTATTAATAACCTGGGTTTTGTG |
| Fw-PdGH32$_{PUL}$ | GCGGCCTGGTGCCGCGCGGCAGCCATATGATGCGAACCGTTAATTTAGT |
| Rev-PdGH32$_{PUL}$ | CGGGCTTTGTTAGCAGCCGGATCTCATTATTTTTCTATTGCTATAT |
| Fw-PdGH68$_{DIS}$ | GTGCCGCGCGGCAGCCATATGATGAATAGTAAAATAGGTAAATCG |
| Rev-PdGH68$_{DIS}$ | CTCGAGTGCGGCCGCAAGCTTCTATTTCTTTAGAGGCTTGATCTGTCC |
| Fw-PdGH32$_{DIS}$-A | GTGCCGCGCGGCAGCCATATGATGTACCGGCGTATCCTCTTCTCCC |
| Rev-PdGH32$_{DIS}$-A | CTCGAGTGCGGCCGCAAGCTTCTAGCCCGCATTACCTTTACTGCG |
| Fw-PdGH32$_{DIS}$-B* | CATCACAGCAGCGGCCTGGTGCCGCGCGGCAGCCATATGACAGAAGAATCATCTA AATACAAGCGCCCAGCTAATTGGGC |
| Rev-PdGH32$_{DIS}$-B* | GTGGTGCTCGAGTGCGGCCGCAAGCTTGTCGACGGAGCTCCTAGCCCGCATTAC CTTTACTGCGAGTGACCGGGGCAACC |

***The distal GH32 (PdGH32$_{DIS}$) was re-cloned to validate an alternative open reading frame (ORF) - using the primer pair marked with a "B" - compared to the originally predicted ORF. To clone the corresponding gene based on the originally predicted ORF, the primer pair marked with an "A" was used. See Table S2 for details.

**Table S2. The predicted and proposed alternative initiator methionine of the non PUL-encoded distal GH32 of *Pseudoalteromonas distincta* (PdGH32$_{DIS}$).**

| | |
|---|---|
| Start codon methionine (bold) of the originally predicted ORF | **M**YRRILFSLVCKWQIKVCSTYSTVSRTVALFFCFTAVISEALHFLILQVFRTAQYHLSCS<br>MSTFAQLHCFVIGAPIRGIMNHNFILIFASRWYMTICVLVVLFCSGMSLAVELGSEAAFS<br>SSPLPLLTEESSKYKRPANWARYRPAVHLTPAKHWMNDPQRPILIDGIWHYYYLYNAD<br>YPIGNGTEWYHATSVDLVHWQDHGVAIDKYKNGLGDIETGSVVIDIDNTAGFGPGAVIA<br>IMTQQHEGVQRQSLFVSTDGGYRFKAYHDNPVMDNPGSEHWRDPKIIWDDTRNEWL<br>MVLAEGHKIGFYTSPDLKHWKYQSGFERDDLGILECPDLFQMSVNGDSATTRWVLAT<br>GANGFQKGMTTGTLYWTGVWDGKNFTVDSDEPQWLDFGADFYAAVTWEDPRHNDA<br>ERLASRYAIGWLNNWAYATKLPTDEWHGGANSIVRRIMLRSVDGKPQLVSQPINAIDK<br>LEGNTVARSNIRVTEASKMALPQPQSDAYRLRVKIDADSDAKEIRFRLKEGNGHFSIVG<br>YNFANDTVFVKRDKDAIANAMPKVYREVRKSPVQARDSFVMLDIIVDTTSIEVFVNNGE<br>VVLSNLVFGAPGANDLSVESIGGDTELSLFELTPLKVAPVTRSKGNAG |
| Proposed alternative start codon methionine (bold) used to re-clone PdGH32$_{DIS}$* (see Table S1) | MYRRILFSLVCKWQIKVCSTYSTVSRTVALFFCFTAVISEALHFLILQVFRTAQYHLSCS<br>MSTFAQLHCFVIGAPIRGIMNHNFILIFASRWYMTICVLVVLFCSGMSLAVELGSEAAFS<br>SSPLPLLTEESSKYKRPANWARYRPAVHLTPAKHWMNDPQRPILIDGIWHYYYLYNAD<br>YPIGNGTEWYHATSVDLVHWQDHGVAIDKYKNGLGDIETGSVVIDIDNTAGFGPGAVIA<br>IMTQQHEGVQRQSLFVSTDGGYRFKAYHDNPVMDNPGSEHWRDPKIIWDDTRNEWL<br>MVLAEGHKIGFYTSPDLKHWKYQSGFERDDLGILECPDLFQMSVNGDSATTRWVLAT<br>GANGFQKGMTTGTLYWTGVWDGKNFTVDSDEPQWLDFGADFYAAVTWEDPRHNDA<br>ERLASRYAIGWLNNWAYATKLPTDEWHGGANSIVRRIMLRSVDGKPQLVSQPINAIDK<br>LEGNTVARSNIRVTEASKMALPQPQSDAYRLRVKIDADSDAKEIRFRLKEGNGHFSIVG<br>YNFANDTVFVKRDKDAIANAMPKVYREVRKSPVQARDSFVMLDIIVDTTSIEVFVNNGE<br>VVLSNLVFGAPGANDLSVESIGGDTELSLFELTPLKVAPVTRSKGNAG |

*predicted signal peptide (blue) using the alternative start codon methionine and the sequence that was cloned (orange) in case of the re-cloned PdGH32$_{DIS}$, which resulted in active CAZyme (see Fig. S13 and S14) compared to inactive CAZyme using the sequence based on the predicted ORF (see Fig. S10)

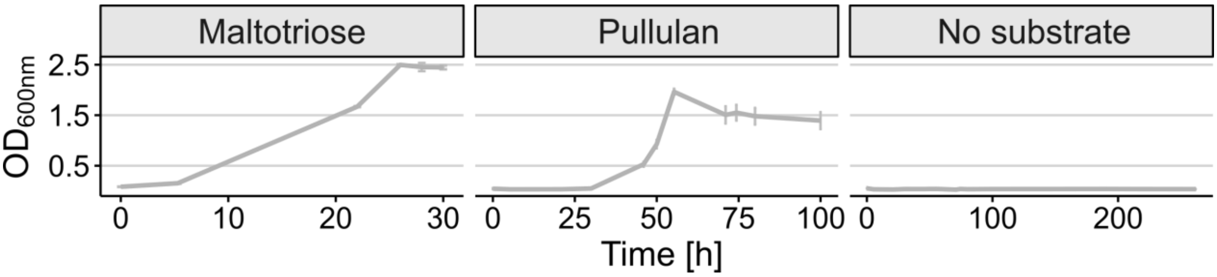


**Fig. S1. Growth controls with α-glucans and no substrate.** Growth of *Pseudoalteromonas distincta* on the α-glucan pullulan and its building block maltotriose, which were used as controls for proteome analysis (whole-cell proteomes, see main text Fig. 2 and Supplementary Dataset 1), and a control without the addition of a carbon source.

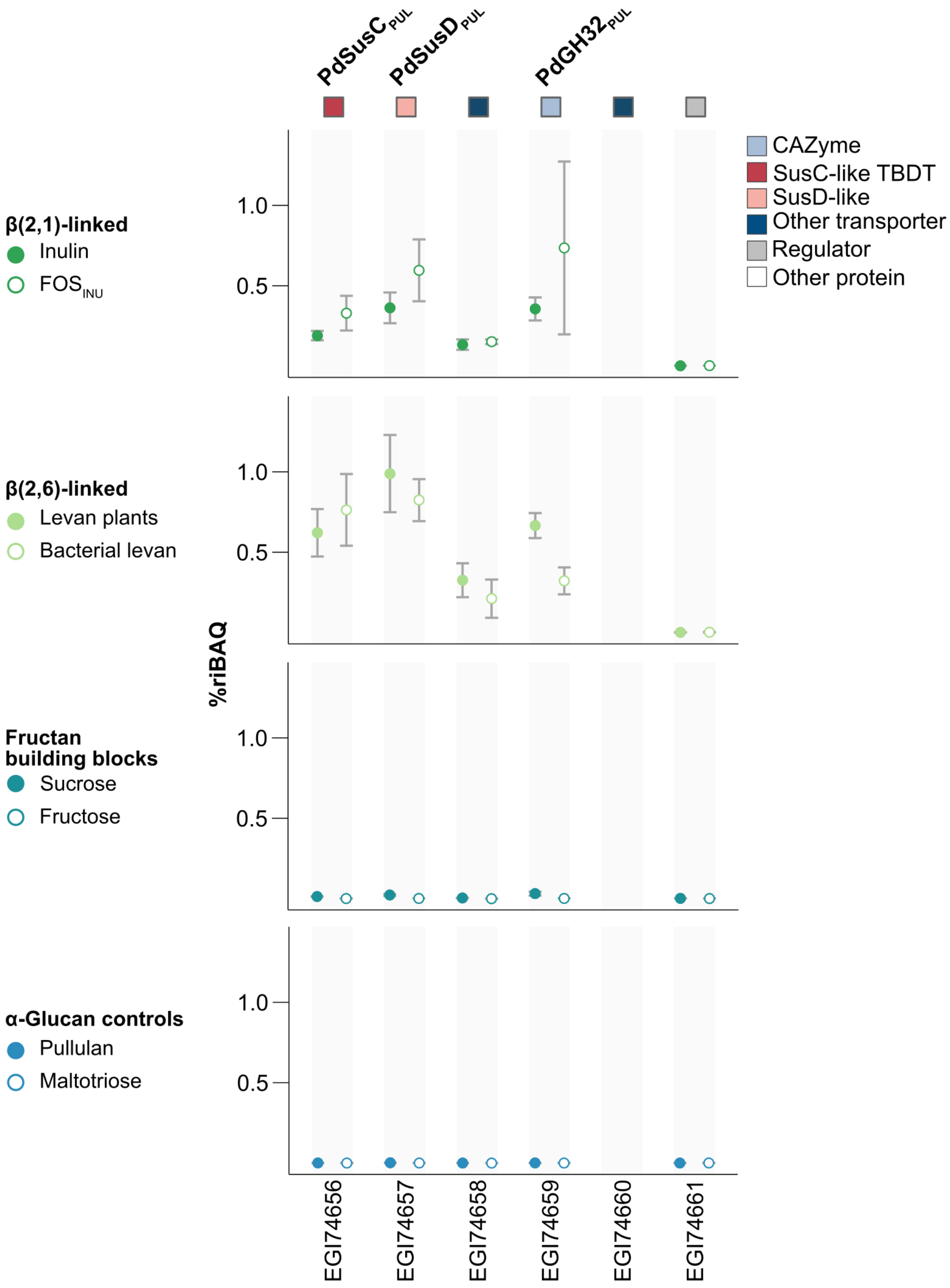


**Fig. S2. Relative protein abundance of fructan PUL-encoded proteins.** Relative protein abundance (%riBAQ) in whole-cell proteomes of *Pseudoalteromonas distincta* grown on different fructose-containing substrates and α-glucans as controls, shown as means (n=3). Individual values and statistics can be found in the Supplementary Dataset 1.

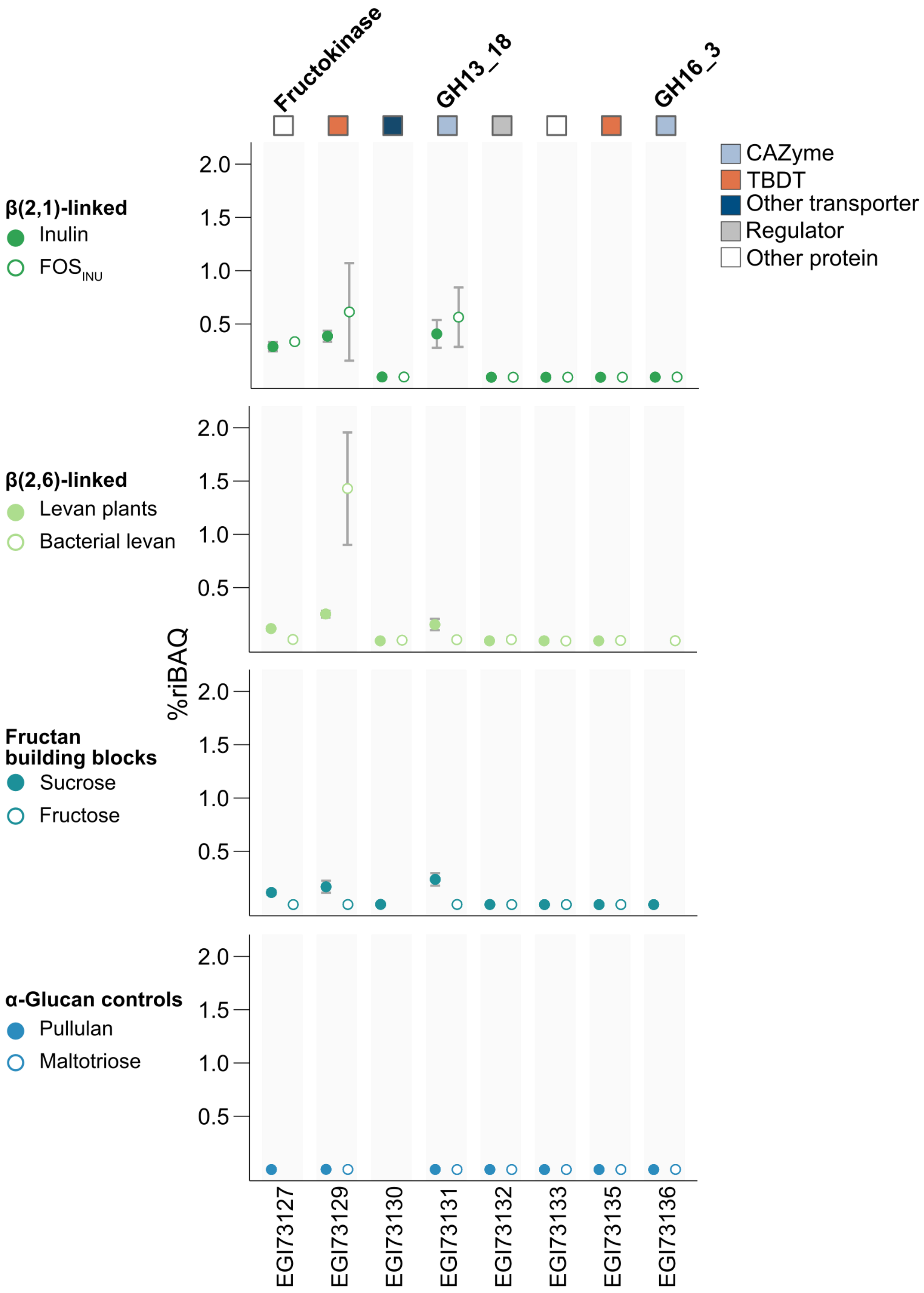


**Fig. S3. Relative abundance of proteins encoded in the sucrose/β-glucan genomic region.** Relative protein abundance (%riBAQ) detected in whole-cell proteomes of *Pseudoalteromonas distincta* grown on different fructose-containing substrates and α-glucans as controls, shown as means (n=3). Individual values and statistics can be found in the Supplementary Dataset 1. EGI73127-33 are associated with sucrose use, while EGI73135 and EGI73136 are presumably associated with β-glucan use.

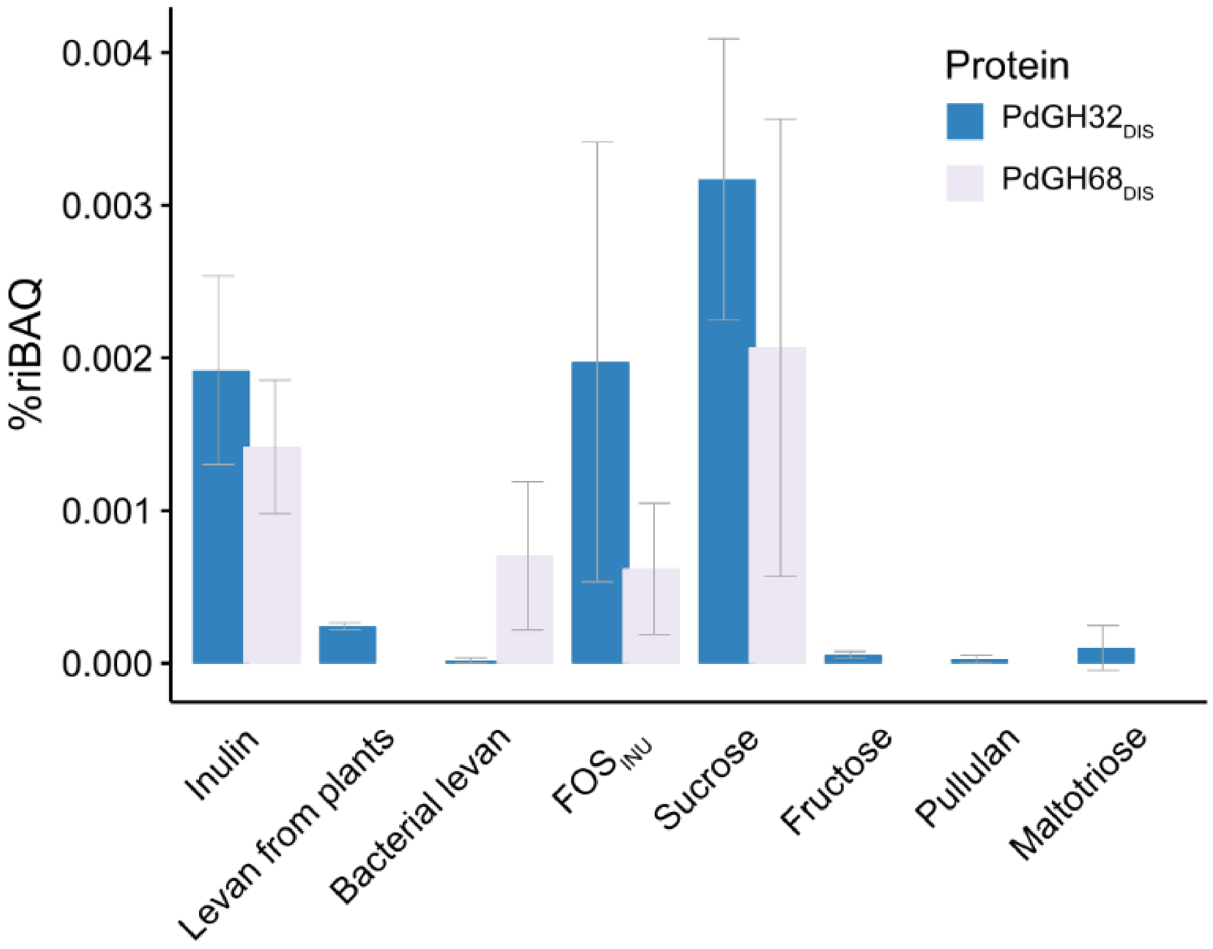


**Fig. S4. Relative protein abundance of the distal PdGH32$_{DIS}$ and PdGH68$_{DIS}$.** Relative protein abundance (%riBAQ) detected in whole-cell proteomes of *Pseudoalteromonas distincta* grown on different fructose-containing substrates and α-glucans as controls, shown as means (n=3). Individual values and statistics can be found in the Supplementary Dataset 1.

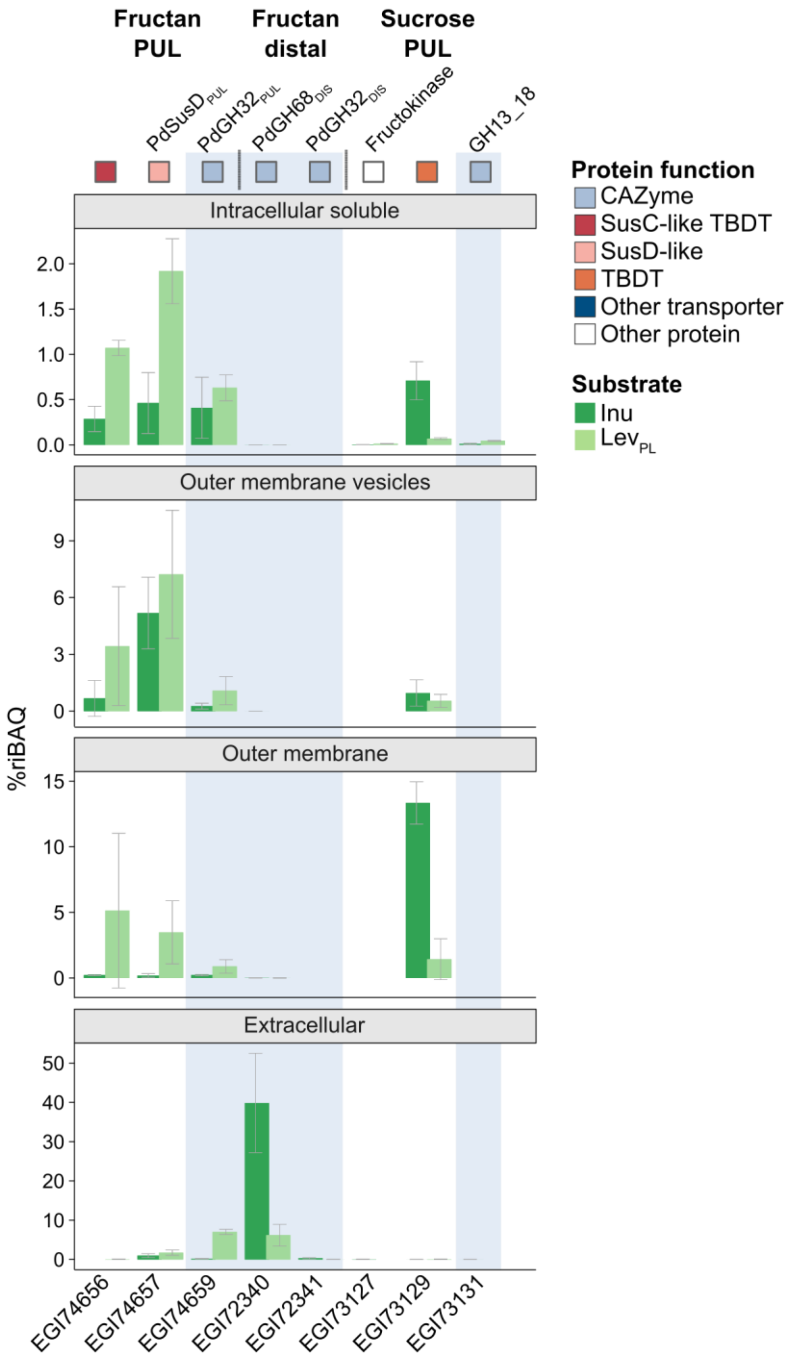


**Fig. S5. The PUL-encoded PdGH32$_{PUL}$ is highly abundant in each analyzed subcellular fraction, while the distal PdGH68$_{DIS}$/PdGH32$_{DIS}$ are associated with the extracellular protein fraction.** Relative abundance (%riBAQ) of selected proteins encoded in the fructan and sucrose PUL as well as of the distal PdGH68$_{DIS}$/PdGH32$_{DIS}$, which are not encoded in a PUL. Relative protein abundance refers to means (n=3) with corresponding standard deviation. Individual values can be found in the Supplementary Dataset 2. Protein abundance should only be compared within the same subcellular fraction due to enrichment bias, the different protocols used and the very different total number of identified proteins per subcellular fraction (e.g., 1672 proteins in the intracellular soluble fraction (inulin) compared to 500 proteins identified in the extracellular protein fraction (inulin); the sum of all protein abundance values (%riBAQs) per sample is 100%). Inu: Inulin, Lev$_{PL}$: Plant-derived levan.

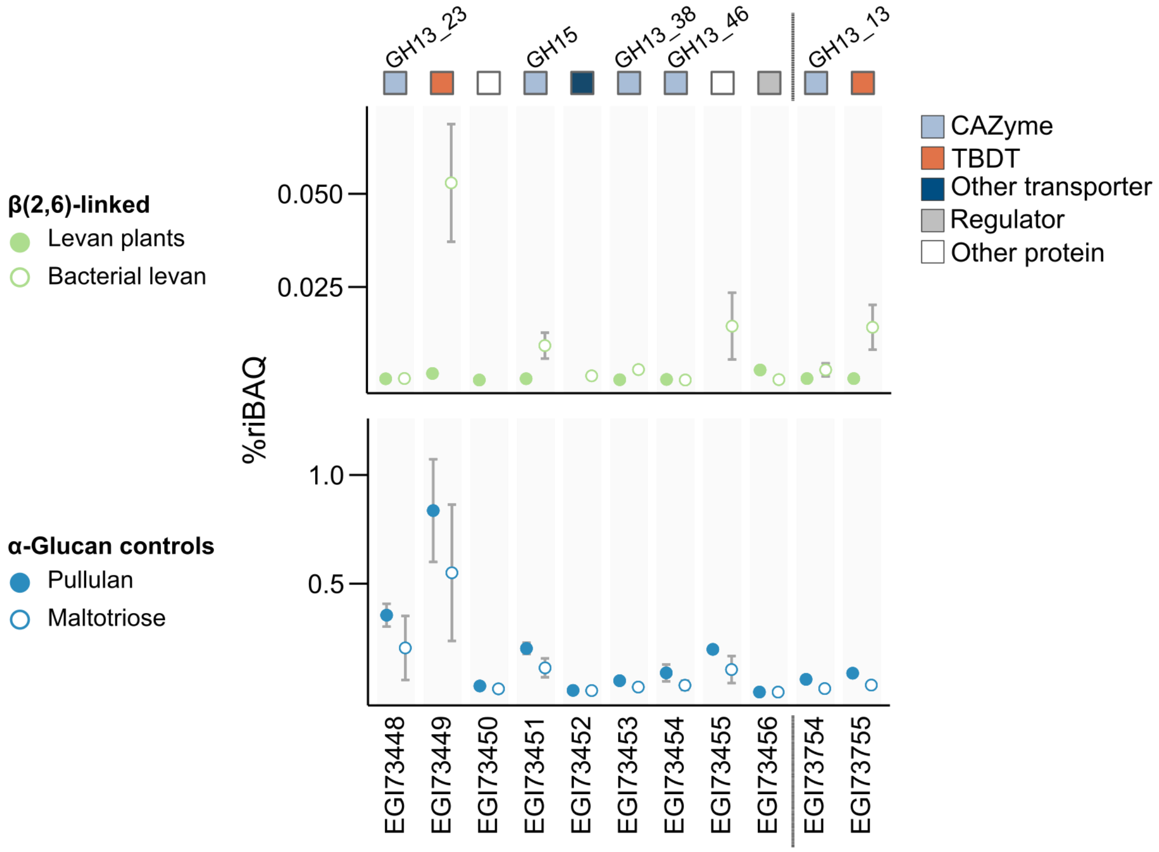


**Fig. S6. Relative protein abundance of α-glucan-related proteins.** Relative protein abundance (%riBAQ) detected in whole-cell proteomes of *Pseudoalteromonas distincta* grown on different fructose-containing substrates and α-glucans as controls, shown as means (n=3). Production refers to α-glucan PUL-encoded proteins (EGI73448-456) and two distal α-glucan-related protein-coding genes (EGI73754/55). There are different scales for the protein abundance values (%riBAQ) for fructan-grown cells and α-glucan-grown cells. Individual values and statistics can be found in the Supplementary Dataset 1.

**Table S3. BlastP results.** The fructan PUL-encoded SusC/D-like proteins of *Pseudoalteromonas distincta* were used as query and sequences from *Bacteroidota* of the human intestine (green) with suspected or known specificity for levan or inulin (1-3) as well as close *Gammaproteobacteria* (blue) and *Bacteroidota* (magenta) homologs from marine, aquatic and soil habitats identified in the phylogenetic analysis (see main text and Fig. S8) were used as subject. Habitats are marked as follows (superscript): A, aquatic; H, human intestine; M, marine; S, soil/sediment.

| PdSusD$_{PUL}$ (EGI74657.1) | | | | | PdSusC$_{PUL}$ (EGI74656.1) | | | | |
|---|---|---|---|---|---|---|---|---|---|
| Subject | Query cover | E value | %Identity | %Positives | Subject | Query cover | E value | %Identity | %Positives |
| A0A290RYI3_9GAMM [M] | 1 | 0 | 99.82 | 100 | A0A290RYI4_9GAMM [M] | 1 | 0 | 97.75 | 98.77 |
| A0A0S1YGD6_9ALTE [M] | 1 | 0 | 77.84 | 86.74 | A0A348MU42_9ALTE [M] | 1 | 0 | 70.12 | 83.6 |
| A0A075NYR5_9ALTE [M] | 1 | 0 | 76.53 | 86.69 | A0AAW7Z595_9ALTE [M] | 1 | 0 | 68.9 | 84.6 |
| A0A348MU43_9ALTE [M] | 1 | 0 | 75.26 | 86.14 | A0A7W2NIU6_9GAMM [M] | 1 | 0 | 67.99 | 84.51 |
| A0A346NLD5_9ALTE [M] | 1 | 0 | 75.31 | 84.24 | A0A075P4A6_9ALTE [M] | 1 | 0 | 66.6 | 82.79 |
| A0A7W2RT51_9GAMM [M] | 1 | 0 | 74.39 | 84.91 | A0A346NLD4_9ALTE [M] | 1 | 0 | 66.63 | 82.4 |
| A0A0X3Y639_9GAMM [S] | 0.99 | 0 | 70.44 | 83.01 | A0A1Y0FTM9_9GAMM [A] | 0.95 | 0 | 64.63 | 79.42 |
| A0A1Y0FTD7_9GAMM [A] | 0.99 | 0 | 69.96 | 82.16 | A0A3N1NME0_9GAMM [M] | 0.98 | 0 | 61.66 | 77.1 |
| A0A3N1NXP7_9GAMM [M] | 1 | 0 | 68.99 | 79.97 | A0A3D4PW26_9GAMM [S] | 0.95 | 0 | 63.45 | 78.03 |
| A0A3D4PW14_9GAMM [S] | 1 | 0 | 68.53 | 80.24 | A0A101M6Y4_9GAMM [S] | 0.98 | 0 | 60.08 | 75.8 |
| A0A2T4X8P2_9BACT [M] | 1 | 0 | 63.68 | 76.84 | I7ZJN8_9GAMM [S] | 0.97 | 0 | 59.5 | 76.01 |
| A0A2I2DNA7_9FLAO [S] | 0.98 | 0 | 61.68 | 75.22 | W7QSA3_9ALTE [M] | 1 | 0 | 56.21 | 72.95 |
| I8TDU5_9GAMM [S] | 0.97 | 0 | 62.5 | 75.18 | A0A2T4X8P5_UNCBA [M] | 0.98 | 0 | 56.44 | 74.95 |
| W7QVY0_9ALTE [M] | 1 | 0 | 53.65 | 69.61 | A0A2I2DN96_9FLAO [S] | 0.96 | 0 | 53.94 | 72.13 |
| A0A534EQF5_9GAMM [S] | 1 | 0 | 52.33 | 66.33 | A0A534HEF4_9GAMM [S] | 0.97 | 0 | 46.87 | 65.33 |
| C9KRX0_9BACE_levan [H] | 1 | 1E-142 | 42.12 | 61.47 | B0NVE1_BACSE_levan [H] | 0.99 | 0 | 43.44 | 61.65 |
| Q8A6W4_BACTN_levan [H] | 0.99 | 7E-129 | 40.92 | 59.08 | Q8A6W3_BACTN_levan [H] | 0.99 | 0 | 43.16 | 62.51 |
| B3CIL3_9BACE_levan [H] | 0.99 | 2E-116 | 39.38 | 57.71 | C9KRX1_9BACE_levan [H] | 0.98 | 0 | 42.8 | 62.51 |
| D1PGX2_9BACT_inulin [H] | 1 | 1E-115 | 38.36 | 56.28 | B3CIL2_9BACE_levan [H] | 0.95 | 0 | 43.41 | 61.54 |
| G6AZL0_9BACT_inulin [H] | 1 | 5E-114 | 37.29 | 54.85 | G6AZL1_9BACT_inulin [H] | 0.98 | 0 | 39.55 | 57 |
| B0NVE2_BACSE_levan [H] | 0.99 | 3E-111 | 39.22 | 56.03 | D1PGX1_9BACT_inulin [H] | 0.98 | 0 | 37.56 | 57 |
| B5D4B2_PHOPM_inulin [H] | 1 | 1E-107 | 37.23 | 53.54 | B5D4B1_PHOPM_inulin [H] | 0.97 | 0 | 37.84 | 56.6 |

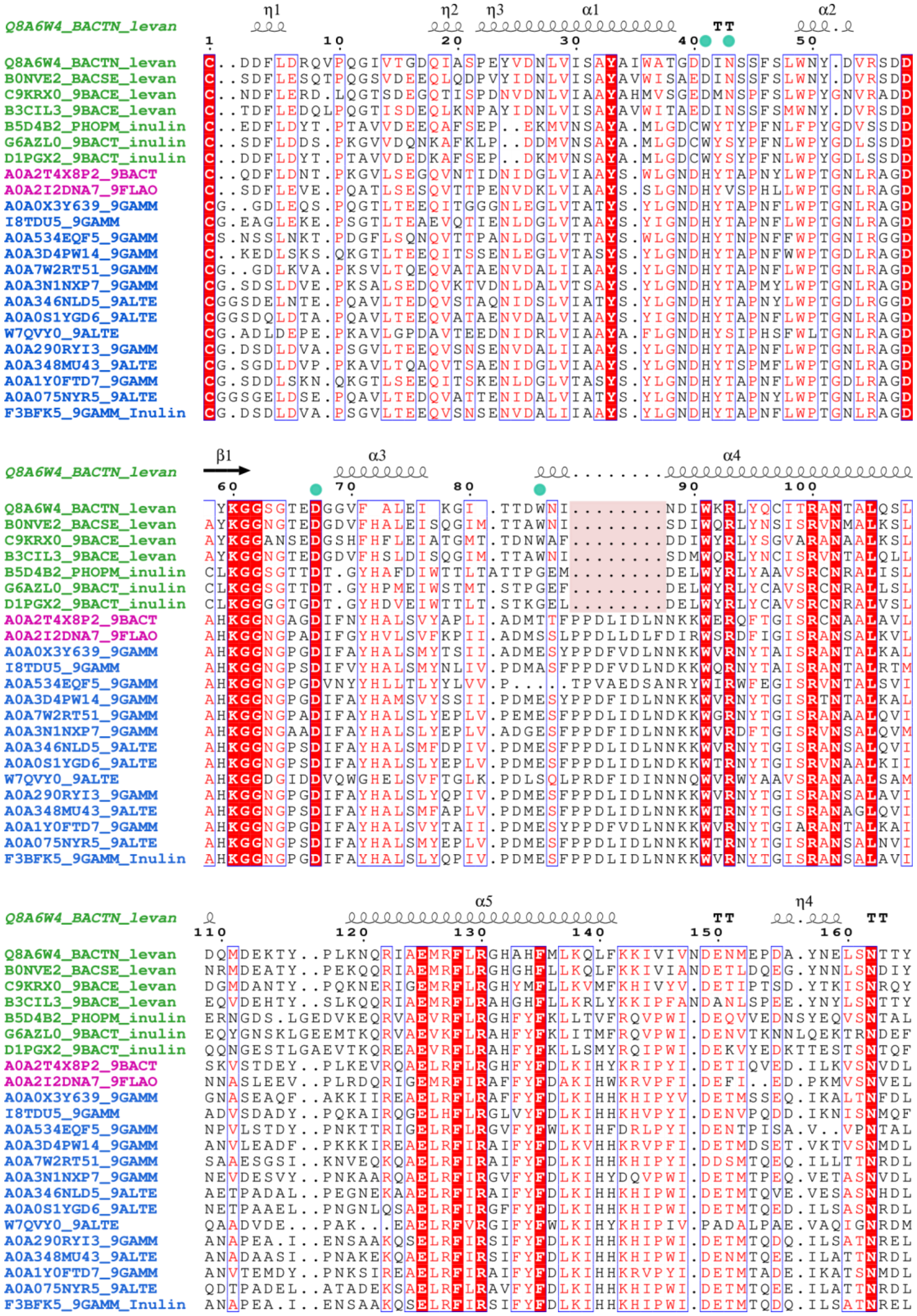
Q8A6W4_BACTN_levan
η1 η2 η3 α1 α2 TT
1 10 20 30 40 50
Q8A6W4_BACTN_levan C..DDFLDRQVPQGIVTGDQIASPEYVDNLVISAYAIWATGDDINSSFSLWNY.DVRSDD
B0NVE2_BACSE_levan C..SDFLESQTPQGSVDEEQLQDPVYIDNLVISAYAVWISAEDINSSFSLWNY.DVRSDD
C9KRX0_9BACE_levan C..NDFLERD.LQGTSDEGQTISPDNVDNLVIAAYAHMVSGEDMNSPFSLWPYGNVRADD
B3CIL3_9BACE_levan C..TDFLEDQLPQGTISDEQLKNPAYIDNLVISAYAVWITAEDINSSFSMWNY.DVRSDD
B5D4B2_PHOPM_inulin C..EDFLDYT.PTAVVDEEQAFSEP..EKMVNSAYA.MLGDCWYTYPFNLFPYGDVSSDD
G6AZL0_9BACT_inulin C..SDFLDDS.PKGVVDQNKAFKLP..DDMVNSAYA.MLGDCWYSYPFNLWPYGDLSSDD
D1PGX2_9BACT_inulin C..DDFLDYT.PTAVVDEDKAFSEP..DKMVNSAYA.MLGDCWYSYPFNLWPYGDLSSDD
A0A2T4X8P2_9BACT C..QDFLDNT.PKGVLSEGQVNTIDNIDGLVIAAYS.WLGNDHYTAPNYLWPTGNLRAGD
A0A2I2DNA7_9FLAO C..SDFLEVE.PQATLSESQVTTPENVDKLVIAAYS.SLGNDHYVSPHLLWPTGNLRAGD
A0A0X3Y639_9GAMM CG..GDLEQS.PQGTLTEEQITGGGNLEGLVTATYS.YLGNDHYTAPNFLWPTGNLRAGD
I8TDU5_9GAMM CG.EAGLEKE.PSGTLTEAEVQTIENLDGLVTAAYS.YIGNDHYTAPNFLWPTGNLRAGD
A0A534EQF5_9GAMM CS.NSSLNKT.PDGFLSQNQVTTPANLDGLVTTAYS.WLGNDHYTNPNFFWPTGNIRGGD
A0A3D4PW14_9GAMM C..KEDLSKS.QKGTLTEEQITSSENLEGLVTASYS.YLGNDHYTAPNFWWPTGDLRAGD
A0A7W2RT51_9GAMM CG..GDLKVA.PKSVLTQEQVATAENVDALIIAAYS.YLGNDHYTAPNFLWPTGNLRAGD
A0A3N1NXP7_9GAMM CG.SDSLDVE.PKSALSEDQVKTVDNLDALVTSAYS.YLGNDHYTAPMYLWPTGNLRAGD
A0A346NLD5_9ALTE CGGSDELNTE.PQAVLTEDQVSTAQNIDSLVIATYS.YLGNDHYTAPNYLWPTGDLRGGD
A0A0S1YGD6_9ALTE CGGSDQLDTA.PQAVLTEEQVATAENVDALVIAAYS.YIGNDHYTAPNFLWPTGNLRAGD
W7QVY0_9ALTE CG.ADLDEPE.PKAVLGPDAVTEEDNIDRLVIAAYA.FLGNDHYSIPHSFWLTGNLRAGD
A0A290RYI3_9GAMM CG.DSDLDVA.PSGVLTEEQVSNSENVDALIIAAYS.YLGNDHYTAPNFLWPTGNLRAGD
A0A348MU43_9ALTE CG.SGDLDVP.PKAVLTQAQVTSAENIDGLVIAAYS.YLGNDHYTSPMFLWPTGNLRAGD
A0A1Y0FTD7_9GAMM CG.SDDLSKN.QKGTLSEEQITSKENLDGLVTASYS.YLGNDHYTAPNFLWPTGDLRAGD
A0A075NYR5_9ALTE CGGSGELDSE.PQAVLTEDQVATTENIDALVIATYS.YLGNDHYTAPNYLWPTGNLRAGD
F3BFK5_9GAMM_Inulin CG.DSDLDVA.PSGVLTEEQVSNSENVDALIIAAYS.YLGNDHYTAPNFLWPTGNLRAGD
Q8A6W4_BACTN_levan
β1 α3 α4
60 70 80 90 100
Q8A6W4_BACTN_levan YKGGSGTEDGGVF ALEI KGI .TTDWNI........NDIWKRLYQCITRANTALQSL
B0NVE2_BACSE_levan AYKGGNGTEDGDVFHALEISQGIM.TTAWNI........SDIWQRLYNSISRVNMALKSL
C9KRX0_9BACE_levan AYKGGANSEDGSHFHFLEIATGMT.TDNWAF........DDIWYRLYSGVARANAALKSL
B3CIL3_9BACE_levan AYKGGNGTEDGDVFHSLDISQGIM.TTAWNI........SDMWQRLYNCISRVNTALQLL
B5D4B2_PHOPM_inulin CLKGGSGTTDT.GYHAFDIWTTLTATTPGEM........DELWYRLYAAVSRCNRALISL
G6AZL0_9BACT_inulin CLKGGSGTTDT.GYHPMEIWSTMT.STPGEF........DELWYRLYCAVSRCNRALLSL
D1PGX2_9BACT_inulin CLKGGGGTGDT.GYHDVEIWTTLT.STKGEL........DELWYRLYCAVSRCNRALVSL
A0A2T4X8P2_9BACT AHKGGNGAGDIFNYHALSVYAPLI.ADMTTFPPDLIDLNNKKWERQFTGISRCNAALAVL
A0A2I2DNA7_9FLAO AHKGGNGPADIFGYHVLSVFKPII.ADMSSLPPDLLDLFDIRWSRDFIGISRANSALKVL
A0A0X3Y639_9GAMM AHKGGNGPGDIFAYHALSMYTSII.ADMESYPPDFVDLNNKKWVRNYTAISRANTALKAL
I8TDU5_9GAMM AHKGGNGPSDIFVYHALSMYNLIV.PDMASFPPDFVDLNDKKWQRNYTAISRANTALRTM
A0A534EQF5_9GAMM AHKGGNGPGDVNYYHLLTLYYLVV.P....TPVAEDSANRYWIRWFEGISRVNTALSVI
A0A3D4PW14_9GAMM AHKGGNGPGDIFAYHAMSVYSSII.PDMESYPPDFIDLNNKKWVRNYTGISRTNTALKAL
A0A7W2RT51_9GAMM AHKGGNGPADIFAYHALSLYEPLV.PEMESFPPDLIDLNDKKWGRNYTGISRANAALQVI
A0A3N1NXP7_9GAMM AHKGGNGAADIFAYHALSLYEPLV.ADGESFPPDFIDLNNKKWVRDFTGISRVNSALQVM
A0A346NLD5_9ALTE AHKGGNGPSDIFAYHALSMFDPIV.PDMESFPPDLIDLNNKKWVRNYTGISRANSALQVI
A0A0S1YGD6_9ALTE AHKGGNGPSDIFAYHALSLYEPLV.PDMESFPPDLIDLNDKKWTRNYTGISRVNAALKII
W7QVY0_9ALTE AHKGGDGIDDVQWGHELSVFTGLK.PDLSQLPRDFIDINNNQWVRWYAAVSRANAALSAM
A0A290RYI3_9GAMM AHKGGNGPGDIFAYHALSLYQPIV.PDMESFPPDLIDLNNKKWVRNYTGISRANSALAVI
A0A348MU43_9ALTE AHKGGNGPSDIFAYHALSMFAPLV.PDMESFPPDLIDLNNKKWTRNYTGISRANAGLQVI
A0A1Y0FTD7_9GAMM AHKGGNGPGDIFAYHALSVYTAII.PDMESYPPDFVDLNNKKWVRNYTGIARANTALKAI
A0A075NYR5_9ALTE AHKGGNGPSDIFAYHALSMYEPLV.PDMESFPPDLIDLNNKKWTRNYTGISRANSALNVI
F3BFK5_9GAMM_Inulin AHKGGNGPGDIFAYHALSLYQPIV.PDMESFPPDLIDLNNKKWVRNYTGISRANSALAVI
Q8A6W4_BACTN_levan
α5 η4 TT TT
110 120 130 140 150 160
Q8A6W4_BACTN_levan DQMDEKTY..PLKNQRIAEMRFLRGHAHFMLKQLFKKIVIVNDENMEPDA.YNELSNTTY
B0NVE2_BACSE_levan NRMDEATY..PEKQQRIAEMRFLRGHGHFLLKQLFKKIVIANDETLDQEG.YNNLSNDIY
C9KRX0_9BACE_levan DGMDANTY..PQKNERIGEMRFLRGHYMFLLKVMFKHIVYV.DETIPTSD.YTKISNRQY
B3CIL3_9BACE_levan EQVDEHTY..SLKQQRIAEMRFLRAHGHFLLKRLYKKIPFANDANLSPEE.YNYLSNTTY
B5D4B2_PHOPM_inulin ERNGDS.LGEDVKEQRVAEVRFLRGHFYFKLLTVFRQVPWI.DEQVVEDNSYEQVSNTAL
G6AZL0_9BACT_inulin EQYGNSKLGEEMTKQRVAEVKFLRAHFYFKLITMFRQVPWI.DENVTKNNLQEKTRNDEF
D1PGX2_9BACT_inulin QQNGESTLGAEVTKQREAEVRFLRAHFYFKLLSMYRQIPWI.DEKVYEDKTTESTSNTQF
A0A2T4X8P2_9BACT SKVSTDEY..PLKEVRQAELRFLRGHFYFDLKIHYKRIPYI.DETIQVED.ILKVSNVDL
A0A2I2DNA7_9FLAO NNASLEEV..PLRDQRIGEMRFIRAFFYFDAKIHWKRVPFI.DEFI..ED.PKMVSNVEL
A0A0X3Y639_9GAMM GNASEAQF..AKKIIREAELRFLRAFFYFDLKIHHKHIPYV.DETMSSEQ.IKALTNFDL
I8TDU5_9GAMM ADVSDADY..PQKAIRQGELHFLRGLVYFDLKIHHKHVPYI.DENVPQDD.IKNISNMQF
A0A534EQF5_9GAMM NPVLSTDY..PNKTTRIGELRFLRGVFYFWLKIHFDRLPYI.DENTPISA.V..VPNTAL
A0A3D4PW14_9GAMM ANVLEADF..PKKKIREAELRFIRAIFYFDLKVHHKRVPFI.DETMDSET.VKTVSNMDL
A0A7W2RT51_9GAMM SAAESGSI..KNVEQKQAELRFIRAIFYFDLKIHHKHIPYI.DDSMTQEQ.ILLTTNRDL
A0A3N1NXP7_9GAMM NEVDESVY..PNKAARQAELRFIRGVFYFDLKIHYDQVPWI.DETMTPEQ.VETASNVDL
A0A346NLD5_9ALTE AETPADAL..PEGNEKAAELRFIRAIFYFDLKIHHKHIPWI.DETMTQVE.VESASNHDL
A0A0S1YGD6_9ALTE NETPAAEL..PNGNLQSAELRFIRGFFYFDLKIHHKHIPWI.DETMTQED.ILSASNRDL
W7QVY0_9ALTE QAADVDE...PAK...EAELRFVRGIFYFWLKIHYKHIPIV.PADALPAE.VAQIGNRDM
A0A290RYI3_9GAMM ANAPEA.I..ENSAAKQSELRFIRSIFYFDLKIHHKHIPWI.DETMTQDQ.ILSATNREL
A0A348MU43_9ALTE ANADAASI..PNAKEKQAELRFIRSIFYFDLKIHHKHIPWI.DENMTQEE.ILATTNRDL
A0A1Y0FTD7_9GAMM ANVTEMDY..PNKSIREAELRFIRAIFYFDLKIHHKRVPYI.DETMTADE.IKATSNMDL
A0A075NYR5_9ALTE QDTPADEL..ATADEKSAELRFLRAVFYFDLKIHHKHIPWI.DETMTQEE.ILATTNRDL
F3BFK5_9GAMM_Inulin ANAPEA.I..ENSAAKQSELRFIRSIFYFDLKIHHKHIPWI.DETMTQDQ.ILSATNREL

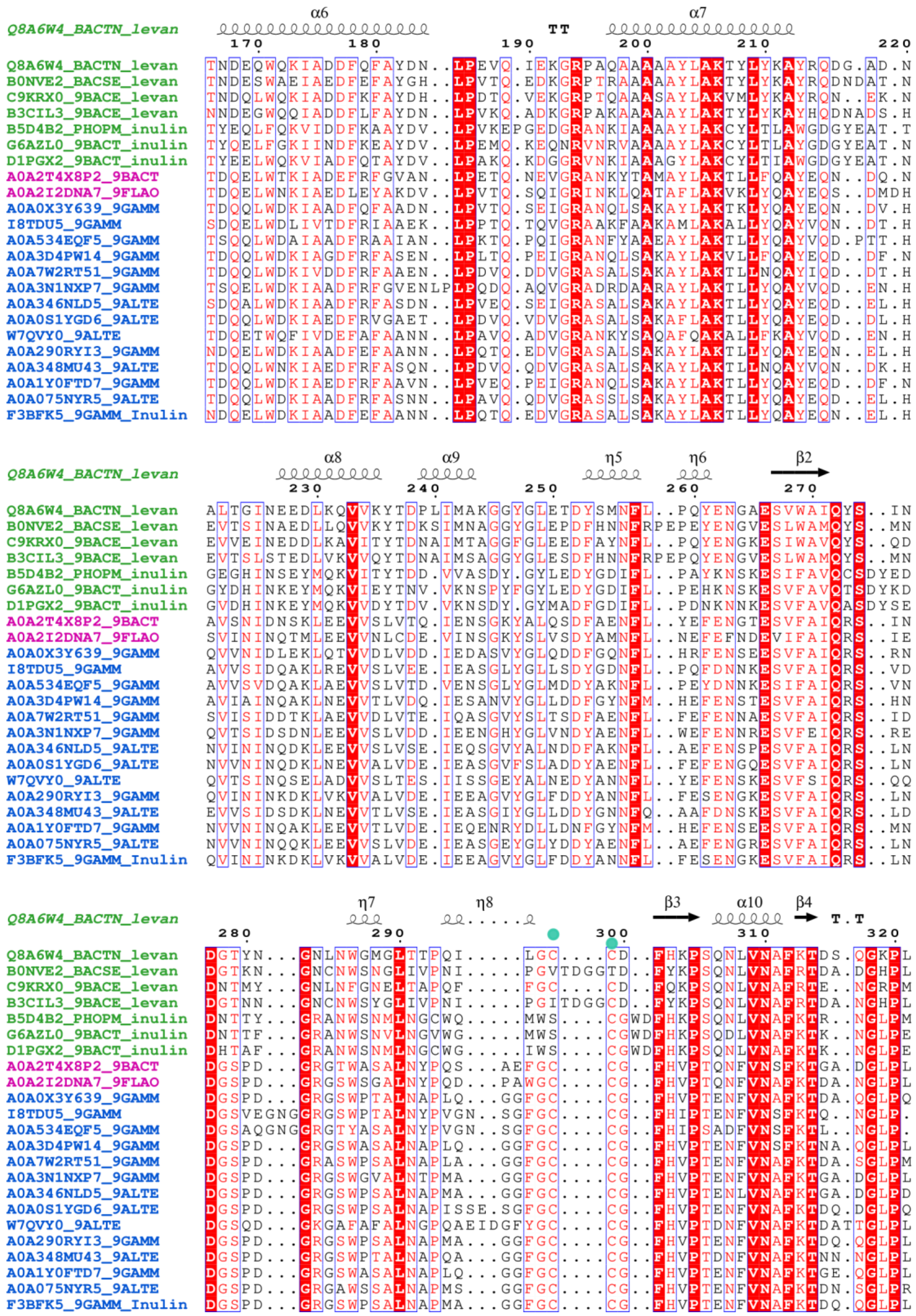
Q8A6W4_BACTN_levan α6 TT α7
170 180 190 200 210 220
Q8A6W4_BACTN_levan TNDEQWQKIADDFQFAYDN..LPEVQ.IEKGRPAQAAAAYLAKTYLYKAYRQDG.AD.N
B0NVE2_BACSE_levan TNDESWAEIAEDFEFAYGH..LPVTQ.EDKGRPTRAAAAYLAKTYLYKAYRQDNDAT.N
C9KRX0_9BACE_levan TNDQLWQKIADDFKFAYDH..LPDTQ.VEKGRPTQAAASAYLAKVMLYKAYRQN..EK.N
B3CIL3_9BACE_levan NNDEGWQQIADDFLFAYDN..LPVKQ.ADKGRPAKAAAAAYLAKTYLYKAYHQDNADS.H
B5D4B2_PHOPM_inulin TYEQLFQKVIDDFKAAYDV..LPVKEPGEDGRANKIAAAAYLAKCYLTLAWGDGYEAT.T
G6AZL0_9BACT_inulin TYQELFGKIINDFKEAYDV..LPEMQ.KEQNRVNRVAAAAYLAKCYLTLAYGDGYEAT.N
D1PGX2_9BACT_inulin TYEELWQKVIADFQTAYDV..LPAKQ.KDGGRVNKIAAAGYLAKCYLTIAWGDGYEAT.N
A0A2T4X8P2_9BACT TDQELWTKIADEFRFGVAN..LPETQ.NEVGRANKYTAMAYLAKTLLFQAYEQN..DQ.N
A0A2I2DNA7_9FLAO TDQELWNKIAEDLEYAKDV..LPVTQ.SQIGRINKLQATAFLAKVKLYQAYEQS..DMDH
A0A0X3Y639_9GAMM TDQQLWDKIAADFQFAADN..LPVTQ.SEIGRANQLSAKAYLAKTKLYQAYEQN..DV.H
I8TDU5_9GAMM SDQELWDLIVTDFRIAAEK..LPPTQ.TQVGRAAKFAAKAMLAKALLYQAYVQN..DT.H
A0A534EQF5_9GAMM TSQQLWDAIAADFRAAIAN..LPKTQ.PQIGRANFYAAEAYLAKTLLYQAYVQD.PTT.H
A0A3D4PW14_9GAMM TDQQLWDKIAGDFRFASEN..LPLTQ.PEIGRANQLSAKAYLAKVLLFQAYEQD..DF.N
A0A7W2RT51_9GAMM TDQQLWDKIVDDFRFAAEN..LPDVQ.DDVGRASALSAKAYLAKTLLNQAYIQD..DT.H
A0A3N1NXP7_9GAMM TSQELWDKIAADFRFGVENLPLPQDQ.AQVGRADRDAARAYLAKTLLYQAYVQD..EN.H
A0A346NLD5_9ALTE SDQALWDKIAADFRFASDN..LPVEQ.SEIGRASALSAKAYLAKTLLYQAYEQD..EL.H
A0A0S1YGD6_9ALTE TDQQLWDKIAEDFRFAAEN..LPDVQ.VDVGRASALSAKAFLAKTLLYQAYEQD..DL.H
W7QVY0_9ALTE TDQETWQFIVDEFAFAANN..LPAVQ.ADVGRANKYSAQAFQAKALLFKAYVQD..EN.H
A0A290RYI3_9GAMM NDQELWDKIAADFEFAANN..LPQTQ.EDVGRASALSAKAYLAKTLLYQAYEQN..EL.H
A0A348MU43_9ALTE TDQQLWNKIAEDFRFASQN..LPDVQ.QDVGRASALSAKAYLAKTLLNQAYVQN..DK.H
A0A1Y0FTD7_9GAMM TDQQLWDKIAEDFRFAAVN..LPVEQ.PEIGRANQLSAKAYLAKTLLYQAYEQD..DF.N
A0A075NYR5_9ALTE TDQQLWDKIAADFRFASNN..LPAVQ.QDVGRASSLSAKAYLAKTLLYQAYEQD..EL.H
F3BFK5_9GAMM_Inulin NDQELWDKIAADFEFAANN..LPQTQ.EDVGRASALSAKAYLAKTLLYQAYEQN..EL.H
Q8A6W4_BACTN_levan α8 α9 η5 η6 β2
230 240 250 260 270
Q8A6W4_BACTN_levan ALTGINEEDLKQVVKYTDPLIMAKGGYGLETDYSMNFL..PQYENGAESVWAIQYS..IN
B0NVE2_BACSE_levan EVTSINAEDLLQVVKYTDKSIMNAGGYGLEPDFHNNFRPEPEYENGVESLWAMQYS..MN
C9KRX0_9BACE_levan EVVEINEDDLKAVITYTDNAIMTAGGFGLEEDFAYNFL..PQYENGKESIWAVQYS..QD
B3CIL3_9BACE_levan EVTSLSTEDLVKVVQYTDNAIMSAGGYGLESDFHNNFRPEPQYENGVESLWAMQYS..MN
B5D4B2_PHOPM_inulin GEGHINSEYMQKVITYTDD.VVASDY.GYLEDYGDIFL..PAYKNSKESIFAVQCSDYED
G6AZL0_9BACT_inulin GYDHINKEYMQKVIEYTNV.VKNSPYFGYLEDYGDIFL..PEHKNSKESVFAVQTSDYKD
D1PGX2_9BACT_inulin GVDHINKEYMQKVVDYTDV.VKNSDY.GYMADFGDIFL..PDNKNNKESVFAVQASDYSE
A0A2T4X8P2_9BACT AVSNIDNSKLEEVVSLVTQ.IENSGKYALQSDFAENFL..YEFENGTESVFAIQRS..IN
A0A2I2DNA7_9FLAO SVININQTMLEEVVNLCDE.VINSGKYSLVSDYAMNFL..NEFEFNDEVIFAIQRS..IE
A0A0X3Y639_9GAMM QVVNIDLEKLQTVVDLVDD.IEDASVYGLQDDFGQNFL..HRFENSEESVFAIQRS..RN
I8TDU5_9GAMM AVVSIDQAKLREVVSLVEE.IEASGLYGLLSDYGDNFL..PQFDNNKESVFAIQRS..VD
A0A534EQF5_9GAMM AVVSVDQAKLAEVVSLVTD.VENSGLYGLMDDYAKNFL..PEYDNNKESIFAIQRS..VN
A0A3D4PW14_9GAMM AVIAINQAKLNEVVTLVDQ.IESANVYGLLDDFGYNFM..HEFENSTESVFAIQRS..HN
A0A7W2RT51_9GAMM SVISIDDTKLAEVVDLVTE.IQASGVYSLTSDFAENFL..FEFENNAESVFAIQRS..ID
A0A3N1NXP7_9GAMM QVTSIDSDNLEEVVSLVDD.IEENGHYGLVNDYAENFL..WEFENNRESVFEIQRS..RE
A0A346NLD5_9ALTE NVININQDKLEEVVSLVSE.IEQSGVYALNDDFAKNFL..AEFENSPESVFAIQRS..LN
A0A0S1YGD6_9ALTE NVVNINQDKLNEVVALVDE.IEASGVFSLADDYAENFL..FEFENGQESVFAIQRS..LN
W7QVY0_9ALTE QVTSINQSELADVVSLTES.IISSGEYALNEDYANNFL..YEFENSKESVFSIQRS..QQ
A0A290RYI3_9GAMM QVININKDKLVKVVALVDE.IEEAGVYGLFDDYANNFL..FESENGKESVFAIQRS..LN
A0A348MU43_9ALTE EVVSIDSDKLNEVVTLVSE.IEASGIYGLLDDYGNNFQ..AAFDNSKESVFAIQRS..LD
A0A1Y0FTD7_9GAMM NVVNINQAKLEEVVTLVDE.IEQENRYDLLDNFGYNFM..HEFENSEESVFAIQRS..MN
A0A075NYR5_9ALTE NVVNINQQKLEEVVSLVDE.IEASGEYGLYDDYAENFL..AEFENGRESVFAIQRS..LN
F3BFK5_9GAMM_Inulin QVININKDKLVKVVALVDE.IEEAGVYGLFDDYANNFL..FESENGKESVFAIQRS..LN
Q8A6W4_BACTN_levan η7 η8 β3 α10 β4 T.T
280 290 300 310 320
Q8A6W4_BACTN_levan DGTYN...GNLNWGMGLTTPQI.....LGC....CD..FHKPSQNLVNAFKTDS.QGKPL
B0NVE2_BACSE_levan DGTKN...GNCNWSNGLIVPNI.....PGVTDGGTD..FYKPSQNLVNAFRTDA.DGHPL
C9KRX0_9BACE_levan DNTMY...GNLNFGNELTAPQF.....FGC....CD..FQKPSQNLVNAFRTE..NGRPM
B3CIL3_9BACE_levan DGTNN...GNCNWSYGLIVPNI.....PGITDGGCD..FYKPSQNLVNAFRTDA.NGHPL
B5D4B2_PHOPM_inulin DNTTY...GRANWSNMLNGCWQ.....MWS....CGWDFHKPSQNLVNAFKTR..NGLPM
G6AZL0_9BACT_inulin DNTTF...GRANWSNVLNGVWG.....MWS....CGWDFHKPSQDLVNAFKTK..NGLPE
D1PGX2_9BACT_inulin DHTAF...GRANWSNMLNGCWG.....IWS....CGWDFHKPSQNLVNAFKTK..NGLPE
A0A2T4X8P2_9BACT DGSPD...GRGTWASALNYPQS...AEFGC....CG..FHVPTQNFVNSFKTGA.DGLPL
A0A2I2DNA7_9FLAO DGSPD...GRGSWSGALNYPQD...PAWGC....CG..FHVPTQNFVNAFKTDA.NGLPL
A0A0X3Y639_9GAMM DGSPD...GRGSWPTALNAPLQ...GGFGC....CG..FHVPTENFVNAFKTDA.QGLPQ
I8TDU5_9GAMM DGSVEGNGGRGSWPTALNYPVGN..SGFGC....CG..FHIPTENFVNSFKTQ..NGLP.
A0A534EQF5_9GAMM DGSAQGNGGRGTYASALNYPVGN..SGFGC....CG..FHIPSADFVNSFKTL..NGLP.
A0A3D4PW14_9GAMM DGSPD...GRGSWASALNAPLQ...GGFGC....CG..FHVPTENFVNSFKTNA.QGLPL
A0A7W2RT51_9GAMM DGSPD...GRASWPSALNAPLA...GGFGC....CG..FHVPTENFVNAFKTDA.SGLPM
A0A3N1NXP7_9GAMM DGSPD...GRGSWGVALNTPMA...GGFGC....CG..FHVPTENFVNAFKTGA.DGLPL
A0A346NLD5_9ALTE DGSPD...GRGSWPSALNAPMA...GGFGC....CG..FHVPTENLVNAFKTGA.DGLPD
A0A0S1YGD6_9ALTE DGSPD...GRGSWPSALNAPISSE.SGFGC....CG..FHVPTENFVNAFKTDQ.DGLPQ
W7QVY0_9ALTE DGSQD...GKGAFAFALNGPQAEIDGFYGC....CG..FHVPTDNFVNAFKTDATTGLPL
A0A290RYI3_9GAMM DGSPD...GRGSWPSALNAPMA...GGFGC....CG..FHVPTENFVNAFKTDQ.QGLPL
A0A348MU43_9ALTE DGSPD...GRGSWPTALNAPQA...GGFGC....CG..FHVPTDNFVNAFKTNN.NGLPL
A0A1Y0FTD7_9GAMM DGSPD...GRGSWASALNAPLQ...GGFGC....CG..FHVPTENFVNAFKTGE.QGLPL
A0A075NYR5_9ALTE DGSPD...GRGAWSSALNAPMS...GGFGC....CG..FHVPTENFVNAFKTDN.SGLPR
F3BFK5_9GAMM_Inulin DGSPD...GRGSWPSALNAPMA...GGFGC....CG..FHVPTENFVNAFKTDQ.QGLPL

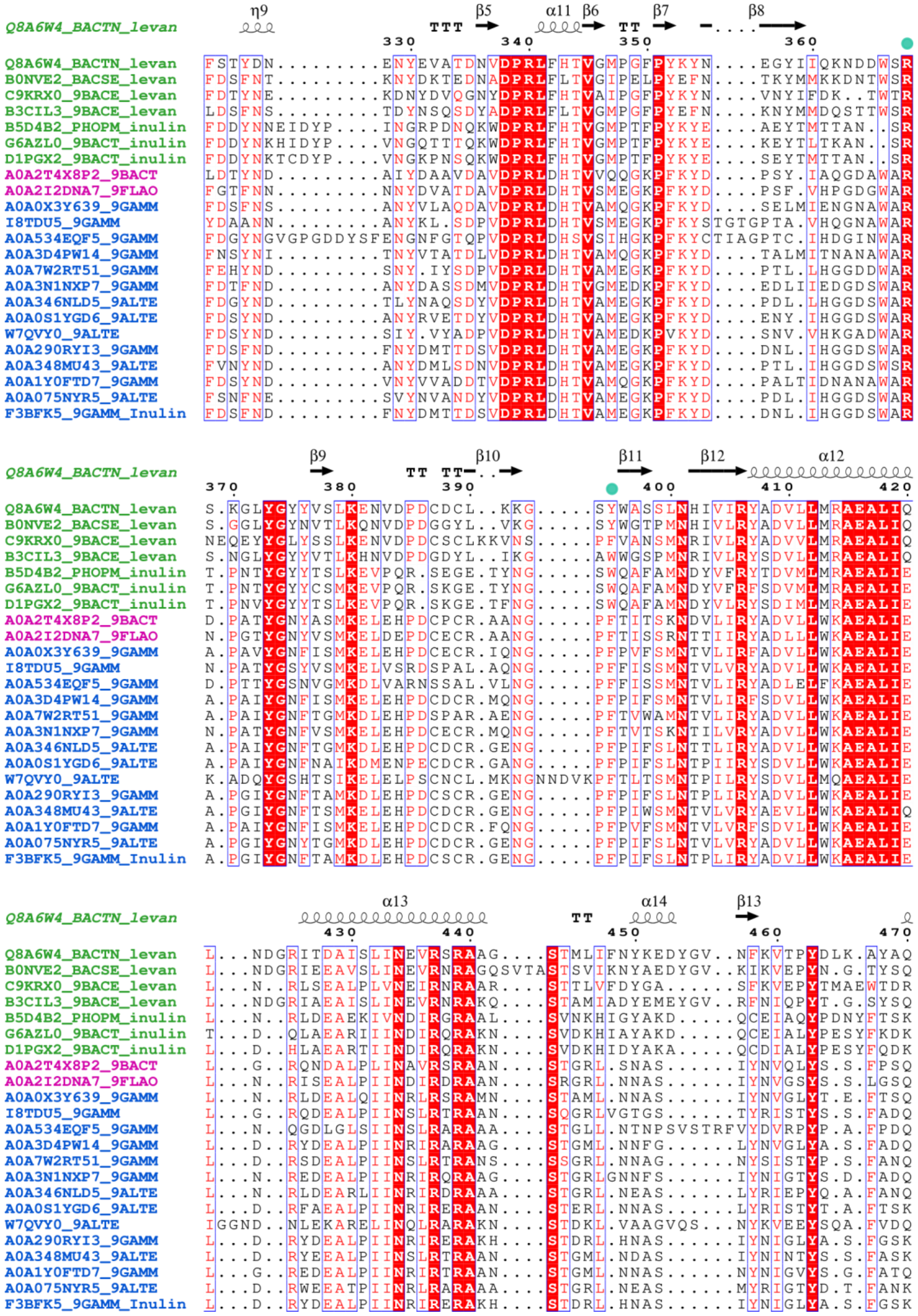
Q8A6W4_BACTN_levan
B0NVE2_BACSE_levan
C9KRX0_9BACE_levan
B3CIL3_9BACE_levan
B5D4B2_PHOPM_inulin
G6AZL0_9BACT_inulin
D1PGX2_9BACT_inulin
A0A2T4X8P2_9BACT
A0A2I2DNA7_9FLAO
A0A0X3Y639_9GAMM
I8TDU5_9GAMM
A0A534EQF5_9GAMM
A0A3D4PW14_9GAMM
A0A7W2RT51_9GAMM
A0A3N1NXP7_9GAMM
A0A346NLD5_9ALTE
A0A0S1YGD6_9ALTE
W7QVY0_9ALTE
A0A290RYI3_9GAMM
A0A348MU43_9ALTE
A0A1Y0FTD7_9GAMM
A0A075NYR5_9ALTE
F3BFK5_9GAMM_Inulin

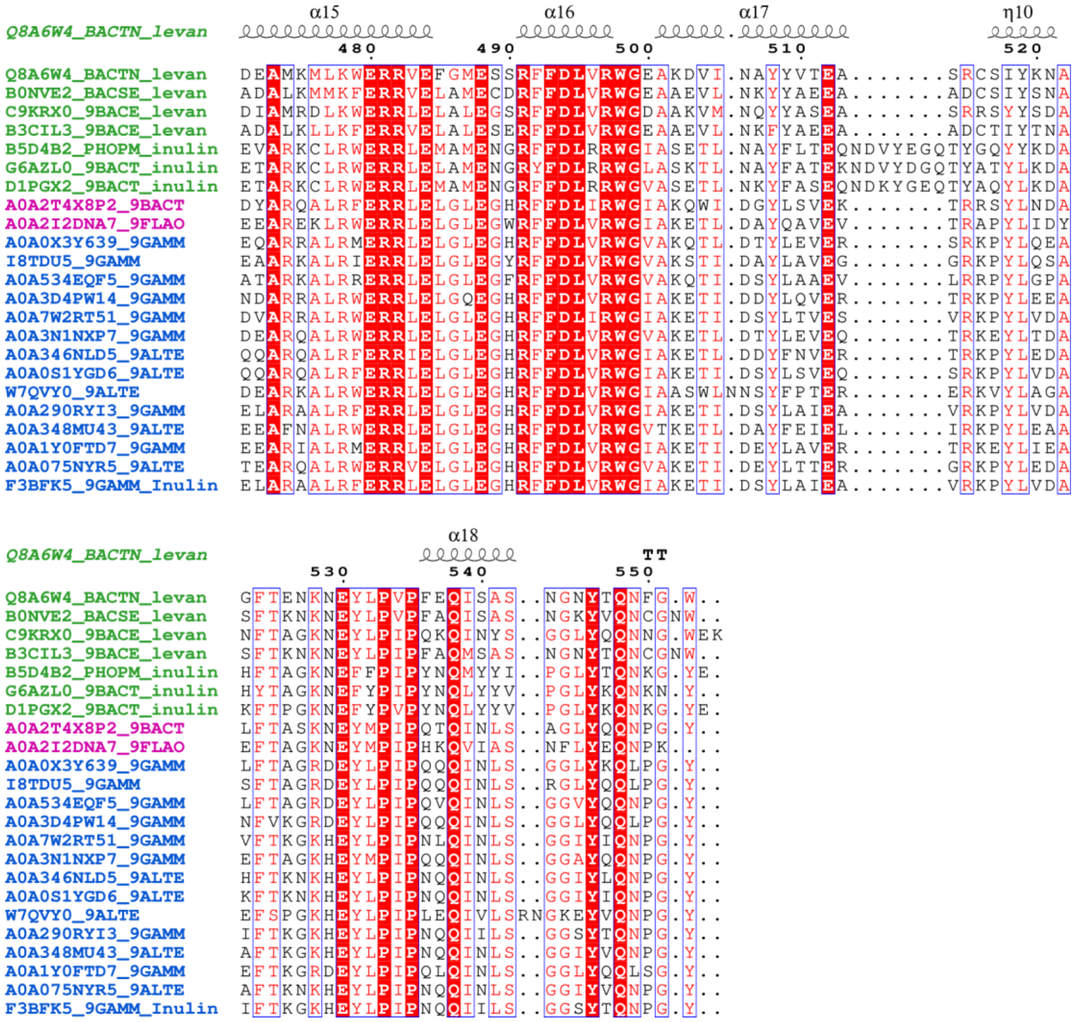

**Fig. S7. Alignment of SusD-like proteins from *Bacteroidota* and *Gammaproteobacteria*.** Displayed is a MAFFT v6.864 alignment (https://www.genome.jp/tools-bin/mafft), using the L-INS-i strategy, of the $FOS_{INU}$-binding $PdSusD_{PUL}$ (Uniprot Identifier F3BFK5_9GAMM) and SusD-like proteins from *Bacteroidota* of the human intestine (green) with suspected or known levan- or inulin-specificity (1-3) as well as close *Gammaproteobacteria* (blue) and *Bacteroidota* (magenta) homologs identified in the phylogenetic analysis (see main text and Fig. S8), which are of marine, aquatic and sediment origin (see Table S3). The Alignment was visualized using ESPript 3.0 (https://espript.ibcp.fr) (4)) and using the 3D crystal structure of the levan-binding SusD-like protein from *Bacteroides thetaiotaomicron* (Uniprot Identifier Q8A6W4_BACTN, PDB ID 5LX8, (1)). Residues involved in binding of levan by the *B. thetaiotaomicron* SusD-like protein (1, 5) are indicated by turquoise circles. A loop (marked rose, preceding α4) in the binding site region is missing in *Bacteroidota* from the human intestine.

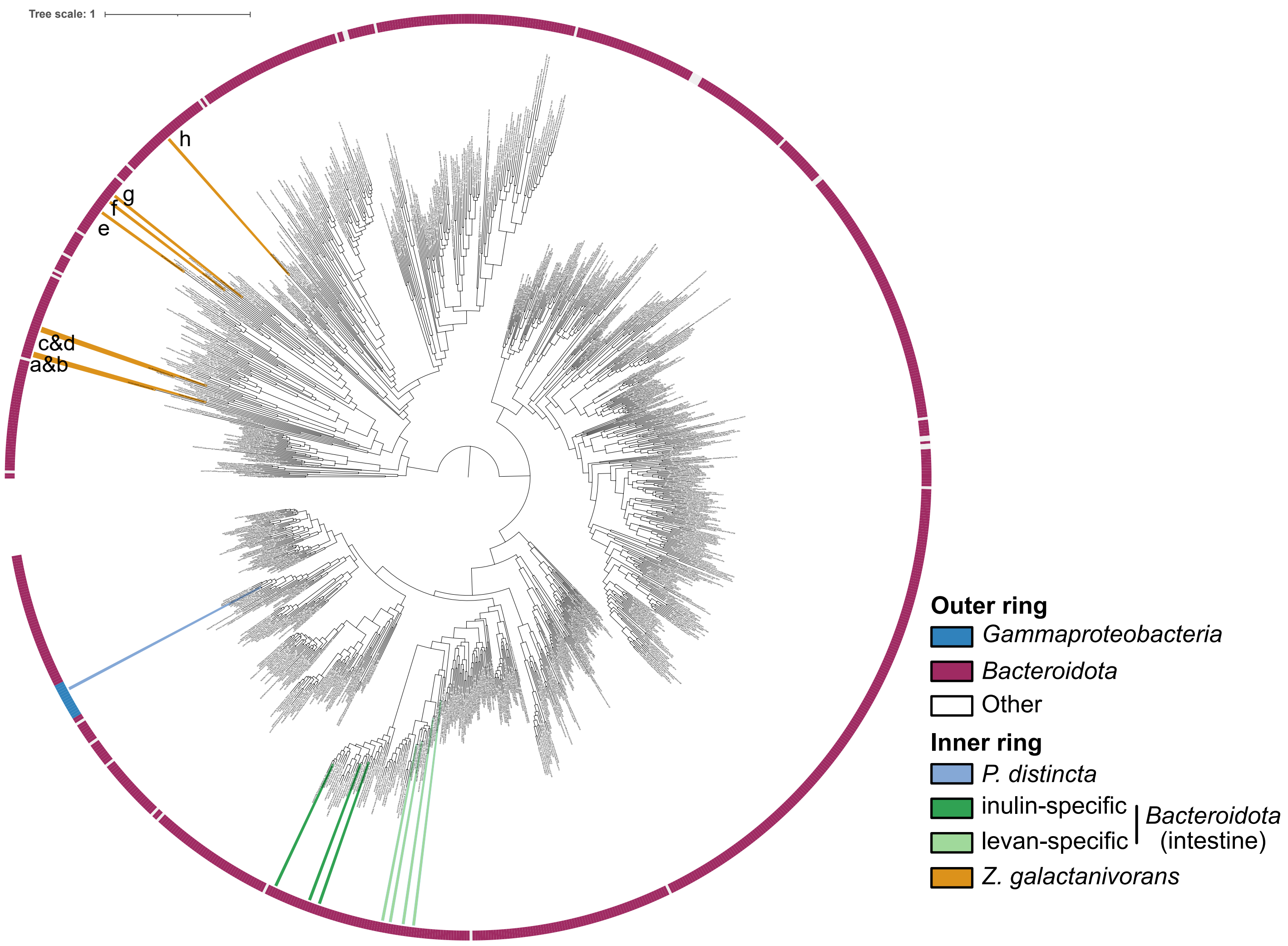


**Fig. S8. Phylogeny of SusD-like sequences.** The outer ring assigns taxonomy, while in the inner ring certain sequences are highlighted, e.g., $PdSusD_{PUL}$ from *Pseudoalteromonas distincta* in light blue (see legend). SusD-like proteins (marked with letters, "a-h") from *Zobellia galactanivorans* $Dsij^T$ (6), which are encoded in PULs targeting polysaccharides other than fructan, were included into the analysis (a: G0L0U6 starch, b: G0L681 alginate, c: G0L189 agar, d: G0L8R1 porphyran, e: G0L194 carrageenan (7), f: G0L886 xylan, g: G0L8P8 unknown sulfated polysaccharide, h: G0L722 unknown sulfated polysaccharide).

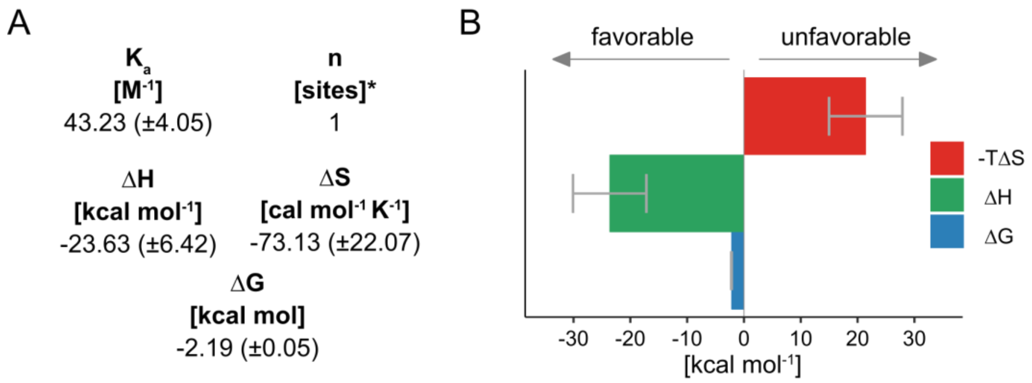


**Fig. S9. Overview of the ITC analysis with PdSusD$_{PUL}$ and FOS$_{INU}$.** (A) Mean values and standard deviation (±) of ITC experiments (n=3). Measurements were performed at 293.15 K. ΔG was calculated using the Gibbs-Helmholtz equation, ΔG=ΔH-TΔS. *The n-value was fixed to 1. (B) Thermodynamics of the interaction visualized as bar graph.

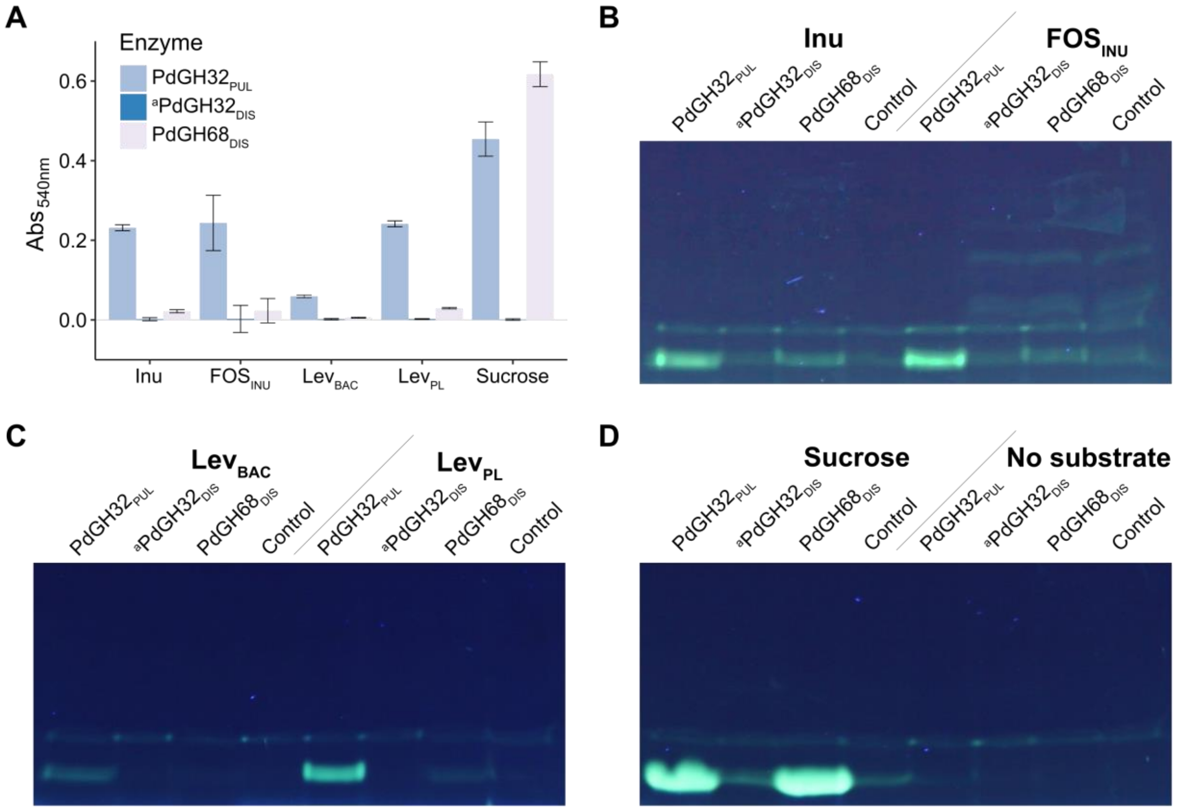


**Fig. S10. Fructan-active CAZymes.** (A) Activity of the PUL-encoded PdGH32$_{PUL}$ and distal PdGH32$_{DIS}$/PdGH68$_{DIS}$ on fructose-containing substrates as determined by RSA (end-point measurements, n=3). Values were corrected against samples without CAZyme. The same samples were used for FACE analyses: (B) β(2,1)-linked substrates, (C) β(2,6)-linked substrates, (D) sucrose. (B-D) In addition, controls without the addition of CAZyme were loaded for each substrate as well as (D) CAZyme controls without the addition of substrate. Inu: Inulin, FOS$_{INU}$: Fructo-oligosaccharides from inulin, Lev$_{BAC}$: Bacterial levan, Lev$_{PL}$: Plant-derived levan. [a]PdGH32$_{DIS}$: protein using the originally predicted initiator methionine (see Table S2), re-cloning led to active enzyme (see Fig. S13 and S14).

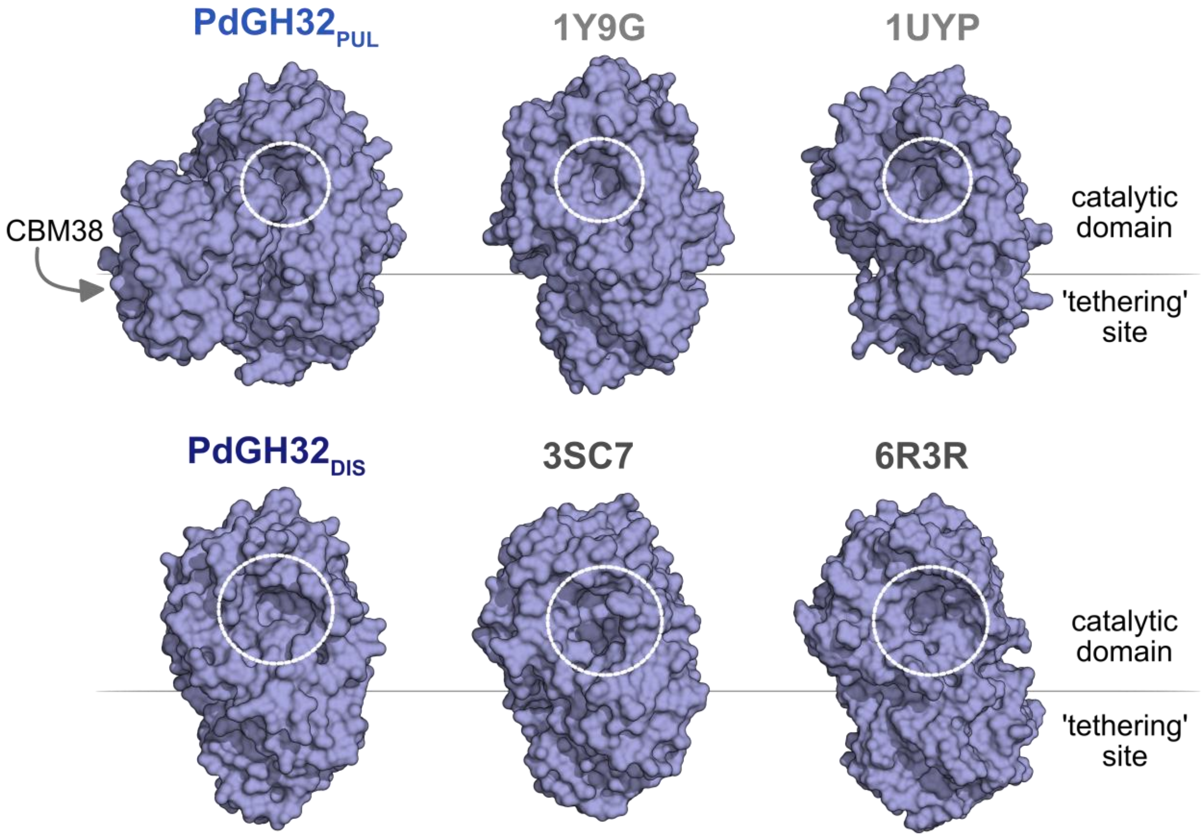


**Fig. S11. Binding grooves of the *Pseudoalteromonas distincta* GH32s.** Surfaces of the PUL-encoded $PdGH32_{PUL}$ and distal $PdGH32_{DIS}$ using AF-predicted structures compared to 3D crystal structures of characterized exo-acting GH32s (1Y9G, *Aspergillus awamori* (8) and 1UYP, *Thermotoga maritima* (9)) and endo-acting GH32s (PDB ID 3SC7, inulinase, *Aspergillus ficuum* (10) and 6R3R, levanase, *Bacteroides thetaiotaomicron* (11)). The catalytic groove is encircled.

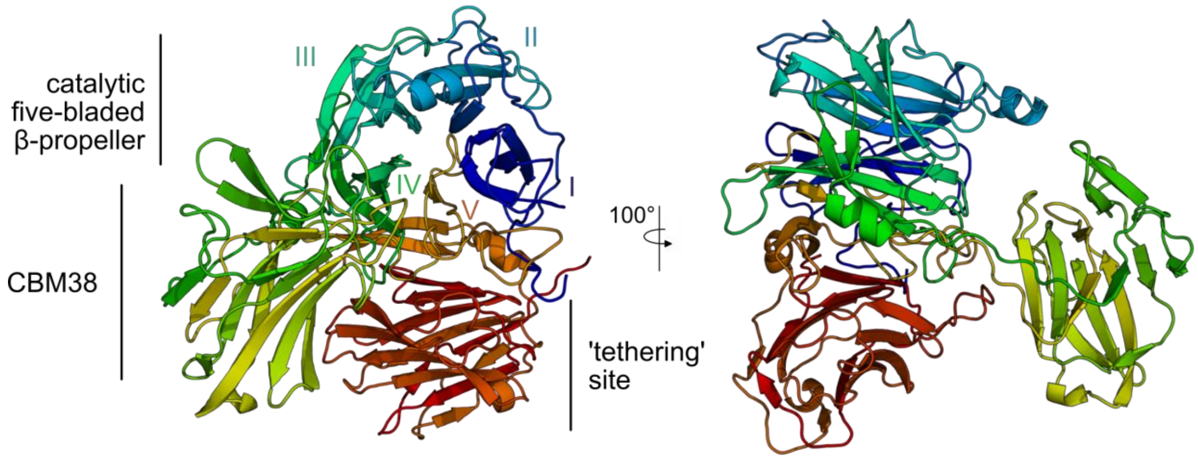


**Fig. S12. A CBM38 in addition to the 'tethering' site in PdGH32$_{PUL}$.** In the case of the PUL-encoded PdGH32$_{PUL}$, a CBM38 protrudes from the five-bladed (I-V) β-propeller catalytic domain, between blade IV and V.

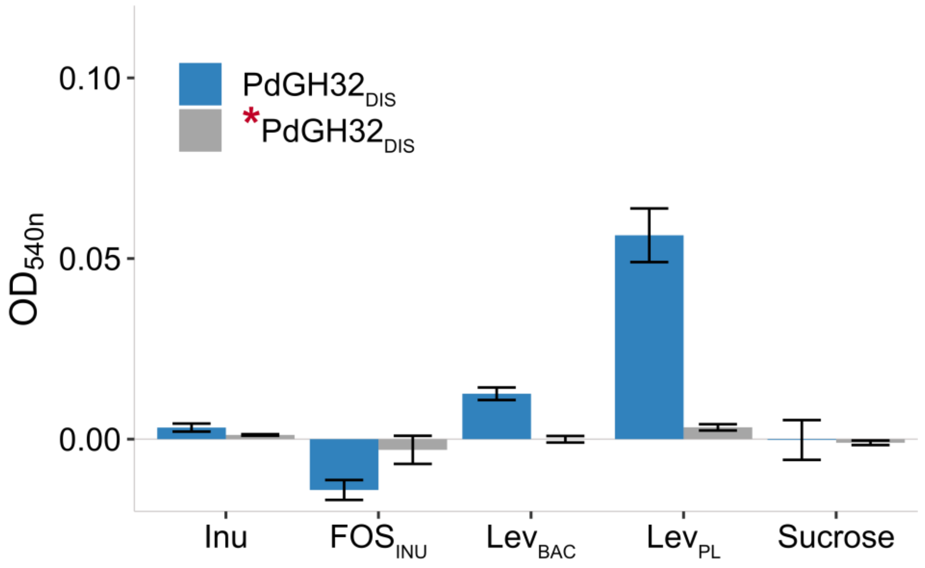


**Fig. S13. The distal PdGH32$_{DIS}$ (re-cloned) is levan-specific.** Activity of the re-cloned PdGH32$_{DIS}$ (see Table S2 for details) compared to heat-inactivated PdGH32$_{DIS}$ (marked with a red asterisk) on different fructans and sucrose as detected by RSA, shown as mean with standard deviation (n=3). Values were corrected against controls without CAZyme. Inu: Inulin, FOS$_{INU}$ Fructo-oligosaccharides from inulin, Lev$_{PL}$: Plant-derived levan, Lev$_{BAC}$: Bacterial levan.

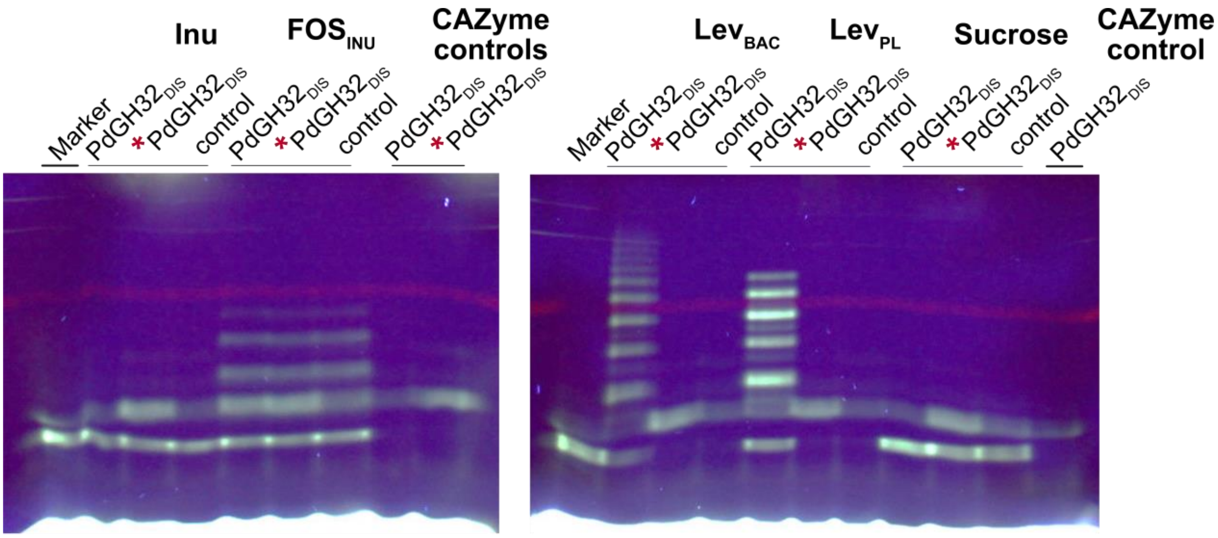


**Fig. S14. The distal PdGH32$_{DIS}$ (re-cloned) is endo-active on levans.** Activity of the re-cloned PdGH32$_{DIS}$ (see Table S2 for details) compared to heat-inactivated PdGH32$_{DIS}$ (marked with a red asterisk) on fructose-containing substrates as determined by FACE analyses. Controls without the addition of CAZyme were loaded for each substrate as well as CAZyme controls without the addition of substrate. The marker contained sucrose and fructose, which could not be separated. Inu: Inulin, FOS$_{INU}$: Fructo-oligosaccharides from inulin (both β(2,1)-linked), Lev$_{BAC}$: Bacterial levan, Lev$_{PL}$: Plant-derived levan (both β(2,6)-linked).

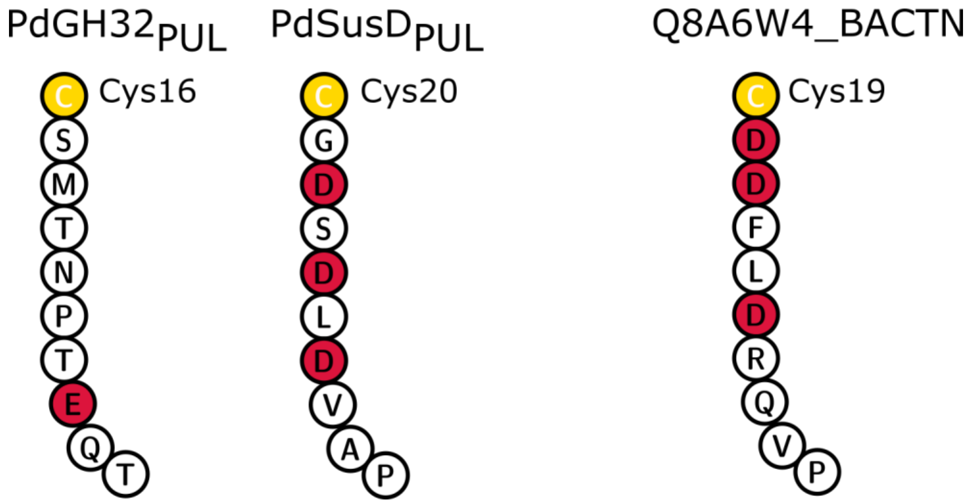


**Fig. S15. Residues following the conserved cysteine in lipoproteins from *Pseudoalteromonas distincta* and *Bacteroides thetaiotaomicron* VPI-5482$^T$.** Shown are 10 amino acids from each protein (*P. distincta*: PdGH32$_{PUL}$ and PdSusD$_{PUL}$, *B. thetaiotaomicron*: Q8A6W4_BACTN, PDB ID 5LX8) starting with the conserved cysteine that is lipidated to the membrane. Acidic residues are highlighted in red. Sequences were visualized using Protter (12).

### References


1. Glenwright AJ, Pothula KR, Bhamidimarri SP, Chorev DS, Baslé A, Firbank SJ, et al. Structural basis for nutrient acquisition by dominant members of the human gut microbiota. Nature. 2017;541(7637):407-11.
2. Joglekar P, Sonnenburg ED, Higginbottom SK, Earle KA, Morland C, Shapiro-Ward S, et al. Genetic Variation of the SusC/SusD Homologs from a Polysaccharide Utilization Locus Underlies Divergent Fructan Specificities and Functional Adaptation in *Bacteroides thetaiotaomicron* Strains. mSphere. 2018;3(3).
3. Sonnenburg ED, Zheng HJ, Joglekar P, Higginbottom SK, Firbank SJ, Bolam DN, et al. Specificity of polysaccharide use in intestinal *Bacteroides* species determines diet-induced microbiota alterations. Cell. 2010;141(7):1241-U256.
4. Robert X, Gouet P. Deciphering key features in protein structures with the new ENDscript server. Nucleic Acids Res. 2014;42(W1):W320-W4.
5. Gray DA, White JBR, Oluwole AO, Rath P, Glenwright AJ, Mazur A, et al. Insights into SusCD-mediated glycan import by a prominent gut symbiont. Nat Commun. 2021;12(1):44.
6. Barbeyron T, Thomas F, Barbe V, Teeling H, Schenowitz C, Dossat C, et al. Habitat and taxon as driving forces of carbohydrate catabolism in marine heterotrophic bacteria: example of the model algae-associated bacterium *Zobellia galactanivorans* Dsij$^{T}$. Environ Microbiol. 2016;18(12):4610-27.
7. Ficko-Blean E, Préchoux A, Thomas F, Rochat T, Larocque R, Zhu YT, et al. Carrageenan catabolism is encoded by a complex regulon in marine heterotrophic bacteria. Nat Commun. 2017;8.
8. Nagem RA, Rojas AL, Golubev AM, Korneeva OS, Eneyskaya EV, Kulminskaya AA, et al. Crystal structure of exo-inulinase from *Aspergillus awamori*: the enzyme fold and structural determinants of substrate recognition. J Mol Biol. 2004;344(2):471-80.
9. Alberto F, Bignon C, Sulzenbacher G, Henrissat B, Czjzek M. The three-dimensional structure of invertase (β-fructosidase) from *Thermotoga maritima* reveals a bimodular arrangement and an evolutionary relationship between retaining and inverting glycosidases. J Biol Chem. 2004;279(18):18903-10.
10. Pouyez J, Mayard A, Vandamme AM, Roussel G, Perpète EA, Wouters J, et al. First crystal structure of an endo-inulinase, INU2, from *Aspergillus ficuum*: discovery of an extra-pocket in the catalytic domain responsible for its endo-activity. Biochimie. 2012;94(11):2423-30.
11. Ernits K, Eek P, Lukk T, Visnapuu T, Alarnäe T. First crystal structure of an endo-levanase - the BT1760 from a human gut commensal *Bacteroides thetaiotaomicron*. Sci Rep. 2019;9.
12. Omasits U, Ahrens CH, Müller S, Wollscheid B. Protter: interactive protein feature visualization and integration with experimental proteomic data. Bioinformatics. 2014;30(6):884-6.
